# NF-κB memory coordinates transcriptional responses to dynamic inflammatory stimuli

**DOI:** 10.1101/2022.01.14.476099

**Authors:** Andrew G. Wang, Minjun Son, Nicholas Thom, Savaş Tay

**Affiliations:** Pritzker School of Molecular Engineering, University of Chicago, Chicago, IL 60637, USA; Medical Scientist Training Program, University of Chicago, Chicago, IL 60637, USA

## Abstract

Many scenarios in cellular communication requires cells to interpret multiple dynamic signals. It is unclear how exposure to immune stimuli alters transcriptional responses to subsequent stimulus under inflammatory conditions. Using high-throughput microfluidic live cell analysis, we systematically profiled the NF-κB response to different signal sequences in single cells. We found that NF-κB dynamics stores the history of signals received by cells: depending on the dose and type of prior pathogenic and cytokine signal, the NF-κB response to subsequent stimuli varied widely, from no response to full activation. Using information theory, we revealed that these stimulus-dependent changes in the NF-κB response encode and reflect information about the identity and dose of the prior stimulus. Small-molecule inhibition, computational modeling, and gene expression profiling show that this encoding is driven by stimulus-dependent engagement of negative feedback modules. These results provide a model for how signal transduction networks process sequences of inflammatory stimuli to coordinate cellular responses in complex dynamic environments.

## Introduction

Exposure to pathogenic stimuli results in acute secretion of inflammatory cytokines, followed by a gradual rise and fall in anti-inflammatory cytokines and growth factors^1–4^. The sequence (temporal ordering) of these stimuli provides information about the local tissue environment to nearby cells, and disruption of this progression is linked to pathology. For example, inflammatory signals in sepsis and chronic inflammation dramatically reshape the innate immune response to subsequent challenges^3,5–7^. Furthermore, efforts to engineer the inflammatory response in adjuvant therapy require understanding how prior exposure alters subsequent stimulus responses^8,9^.

Despite the diversity of inflammatory signals, many of these converge on few signaling networks with shared intracellular kinases and activated transcription factors. For example, pathogenic ligands which activate the Toll-like receptor (TLR) family and pro-inflammatory cytokines secreted by host macrophages all converge on a small set of key inflammatory transcription factors, including the canonical NF-κB family transcription factor RelA^10–12^. Patterns of NF-κB activation over time, or activation dynamics, transmit information about stimulus identity and coordinate the subsequent inflammatory response. Ligands induce distinct dynamics of NF-κB nuclear translocation, which facilitate accurate information transmission from extracellular signals to expression of response genes^13,14^. NF-κB dynamics reshape the epigenetic landscape of the cell and regulate gene expression induced by each stimulus^15,16^. However, it is unknown how prior signal exposure alters NF-κB dynamics. If information about prior stimuli is stored through changing intracellular signaling networks, it raises the possibility that activation dynamics can reflect both the cell’s current stimulus and prior stimulus history.

Previous studies of innate immune signaling focused on population-level effects of stimulus history at timescales of days to weeks^3,5,17,18^. These studies report that innate immune memory can induce both priming, where response to subsequent stimulus is stronger^6,18^, and tolerance, where the subsequent response becomes attenuated^5,19,20^. However, innate immune memory at short timescales is poorly studied due to the difficulties in strict control of stimulus timing and continual cell monitoring. Furthermore, population averaged read-outs often blur single cell dynamics and may not represent the actual cellular response.

Here, we explored how prior stimulus history alters subsequent signaling responses in the NF-κB signaling network by combining automated microfluidic stimulation with live cell imaging (Fig 1A). We found that prior stimuli produced distinct attenuation patterns in subsequent NF-κB signaling dynamics through differential regulation of negative feedbacks. These patterns encode information about the cell’s prior history, showing that the NF-κB network stores information about the temporal sequence of environmental signals and transmits that information in the inflammatory response.

**Figure 1:**
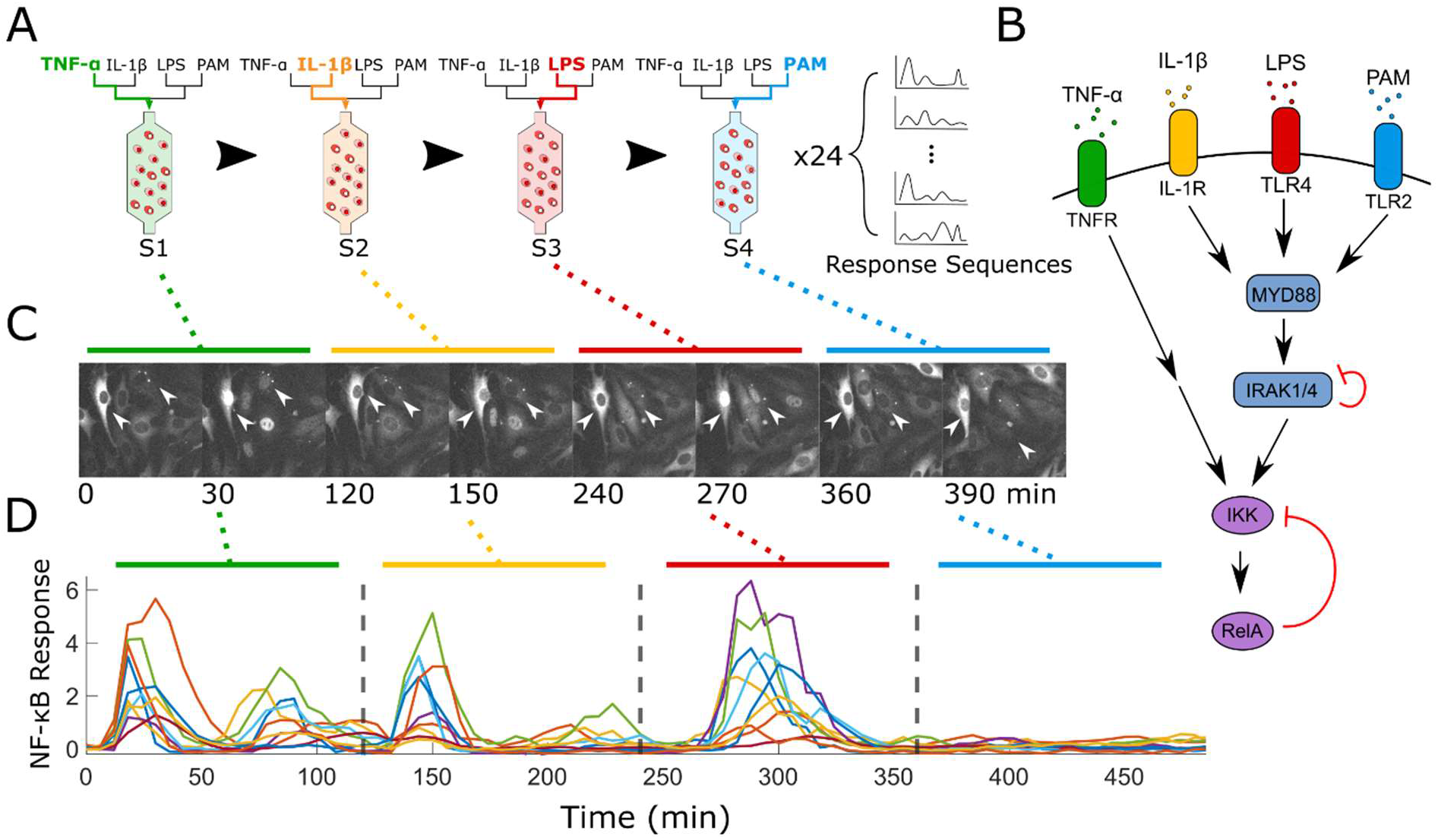
Microfluidic live cell imaging tracks single cell NF-κB responses through multiple sequential stimuli. A) Schematic representation of experimental conditions and microfluidic imaging set up. All non-repeating combinations of 4 ligands were provided to RelA-DsRed tagged RelA^-/-^ 3T3s grown in an automated microfluidic cell culture device and response dynamics were measured. B) Schematic representation of TNF-α (TNFR), IL-1β (IL-1R), LPS (TLR4), and PAM (TLR2) signaling converging on activation of RelA. C) Representative grayscale images of RelA nuclear translocation over 480 minutes of stimulation with mid dose TNF-α (0 min), IL-1β (120 min), LPS (240 min), and PAM (360 min). RelA nuclear translocation into the nucleus in single cells (white arrows) is shown depending on the supplied ligand. D) Quantification of nuclear/cytoplasmic NF-κB over imaging interval for the same condition as in C). Gray dashed lines indicate when new stimulus was provided.

## Results

### Prior ligand history influences NF-κB activation to subsequent stimuli

We focused on the interactions between four inflammatory ligands, tumor necrosis factor alpha (TNF-α), interleukin 1β (IL-1β), lipopolysaccharide (LPS), and PAM2CSK4 (PAM). TNF-α and IL-1β are key pro-inflammatory cytokines which are secreted by sentinel cells and which activate TNFR and IL1R respectively^21^. LPS is a cell wall component of Gram-negative bacteria which activates TLR4, while PAM is a synthetic analogue of bacterial lipopeptides which activates TLR2/6^10^. Thus, LPS and PAM represent pathogen signals, which would trigger local secretion of TNF-α and IL-1β in an infection scenario. Signaling for LPS, PAM, and IL-1β share the receptor-associated adaptor protein MyD88 and downstream components, including IRAK1 (Fig 1B)^22^. In contrast, TNF-α signaling acts through a different set of receptor-associated intermediaries^11^. All these pathways converge at activation of IκB-kinase (IKK), which mediates nuclear translocation of RelA^10,11^. Multiple levels of negative feedback regulate this network, including autoinhibitory phosphorylation of IRAK1 and several transcriptionally regulated negative feedback proteins, such as A20 and IκB∊ (Fig 1B)^23–26^. Each of these negative feedback proteins targets different components in the NF-κB signaling network (Fig 1B)^23^.

To characterize how prior histories shape the NF-κB response to a subsequent ligand, we used a microfluidic platform to provide sequential stimuli to RelA^-/-^ NIH/3T3 fibroblasts (3T3s) expressing a RelA-DsRed fusion protein (Fig 1A)^24,27^. By continuously imaging 3T3s in this platform, we evaluated NF-κB dynamics under a series of stimuli without disrupting the cells (Fig 1C-D). We first systematically profiled the effects of prior history by stimulating cells with non-repeating sequences of all four ligands. This approach produced 24 unique stimulus conditions. The first stimulus (S1) is provided to cells without prior inflammatory ligand exposure, and thus induces a “naïve” response. However, the second, third and fourth stimuli (S2-4) would induce NF-κB responses affected by one, two, or three prior ligands, respectively. We used the response to a particular ligand at S1 as a baseline for comparing how different prior stimulus sequences change the response to that ligand. Additionally, to test how stimulus dose changes prior history effects, we calibrated high, mid, and low doses for each ligand based on the percentage of activated cells (Supp. Fig 1), then repeated the 24 stimulus sequences for each dose. In our initial dataset of 72 conditions, we analyzed more than 10,000 single cells (Fig 2A-C, Supp. Fig 2-4) with a range of prior histories and stimulus doses.

**Figure 2:**
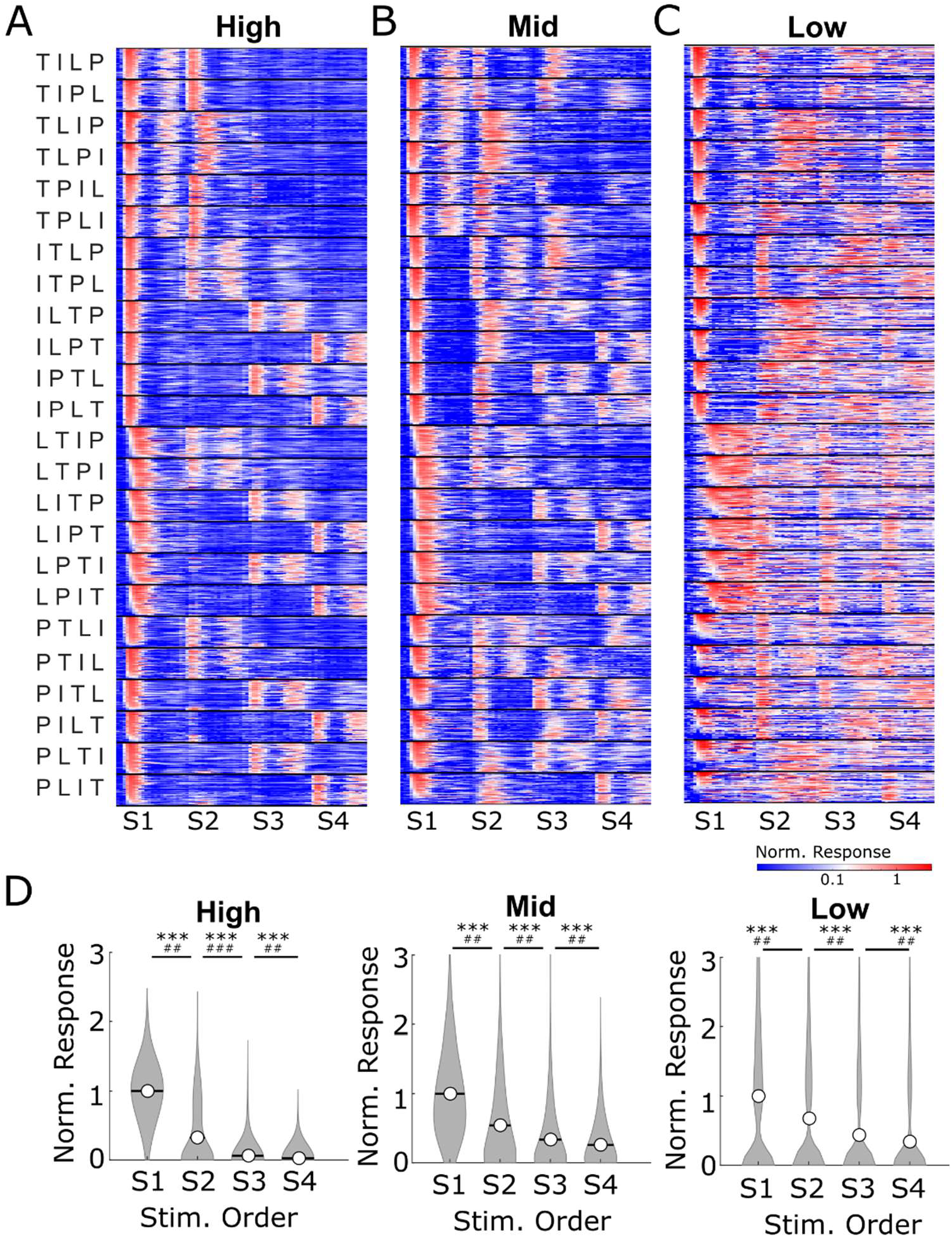
Single-cell NF-κB activation traces reveal ligand and dose specific attenuation of signaling by prior stimuli. A-C) NF-κB response dynamics over 2 hours of stimulus for each ligand normalized to the peak amplitude of the mean response at S1 (naïve). 50 randomly selected single-cell traces are displayed for each condition. Each row shows the nuclear NF-κB level of a single-cell measured by time-lapse microscopy, and x-axis shows the time. Heatmap columns are arranged from the first stimulus (S1) to the fourth stimulus (S4). Stimulus orders are shown to the left of the first heatmap, where T stands for TNF-α, I for IL-1β, L for LPS, and P for PAM. D) Single cell responses from S1-S4, normalized to the mean of corresponding S1 response (>2000 cells for each condition). Open circle and line show the mean. Bonferroni corrected Wilcoxon rank sum test p-value < 10^−4^ (***). Fold change difference between sample means > 1 (#), > 1.25 (##), or >4 (###).

To observe general trends in ligand response, we first examined how the response to a specific ligand changed depending on its order in a stimulus sequence. All single cell responses in each sequence position were grouped by ligand and normalized to the mean S1 response for that ligand (Fig 2A-C, Supp. Fig 5). When we compared the amplitude changes over the four sequence positions, we observed that response for each ligand decreased from S1 to S4 (Fig 2D, Supp. Fig 6). Even in low dose conditions, where response heterogeneity results in highly variable response amplitudes, ligand responses decreased from S1 to S4. Thus, we concluded that prior exposure history primarily attenuates signaling responses to subsequent ligands. However, we also noted that distinct patterns of attenuation existed depending on ligand identity and dose. Even at high dose, where attenuation was strongest, cells responded to TNF-α stimulus irrespective of prior history (Fig 2A, Supp. Fig 5A). At mid and low dose, each ligand displayed different history responses. LPS and TNF-α responses exhibited the weakest attenuation, with some level of stimulus response retained across most conditions, while IL-1β and PAM responses showed large variability in response depending on prior stimulus history (Fig 2B-C, Supp. Fig 5B-C). Thus, particular ligand histories can alter subsequent responses in a consistent and predictable manner.

### The NF-κB network reflects information about prior ligands in the subsequent response

If particular ligand histories alter subsequent response dynamics in a distinctive manner, it would be possible to characterize a cell’s prior history through its response to subsequent stimuli. However, the regulation of a genetic network is inherently noisy, resulting in diverse response to identical stimulus at the single cell level and over time ^28–30^. This variability may impact how accurately individual cells can reflect prior history in subsequent responses. Thus, we needed to address single cell variability in characterizing how effectively prior history is reflected in subsequent response.

We used information theory to characterize the distinguishability of NF-κB responses to different stimulus orders despite single cell noise. In information theory, the maximum information transmittable by a noisy network is described by the channel capacity (CC) (Fig 3A). In our case, the CC represents the maximum distinguishability of groups in a population response. Therefore, the CC can be used to quantify the accuracy of signal transduction in the NF-κB network^13,31–33^. We first measured the CC of the NF-κB network in distinguishing all 24 stimulus conditions in each dose. If the NF-κB network did not retain information about prior history, we would expect the CC to stay the same or decrease from S1 to S4, since the effect of noise is enhanced with signal attenuation (Fig 2D)^34^. However, we found that CC increased from S1 to S2 despite attenuation (Fig 3B). Even later in the stimulus sequence at S3 and S4, where attenuation became more pronounced, the CC still remained above the baseline at S1. These observations indicate that, even though the same four ligands are used for stimulation in each sequence, more distinguishable responses are present in S2 – 4. Thus, the NF-κB signaling network retains information about prior history and coordinates subsequent stimulus responses based on prior exposure.

**Figure 3:**
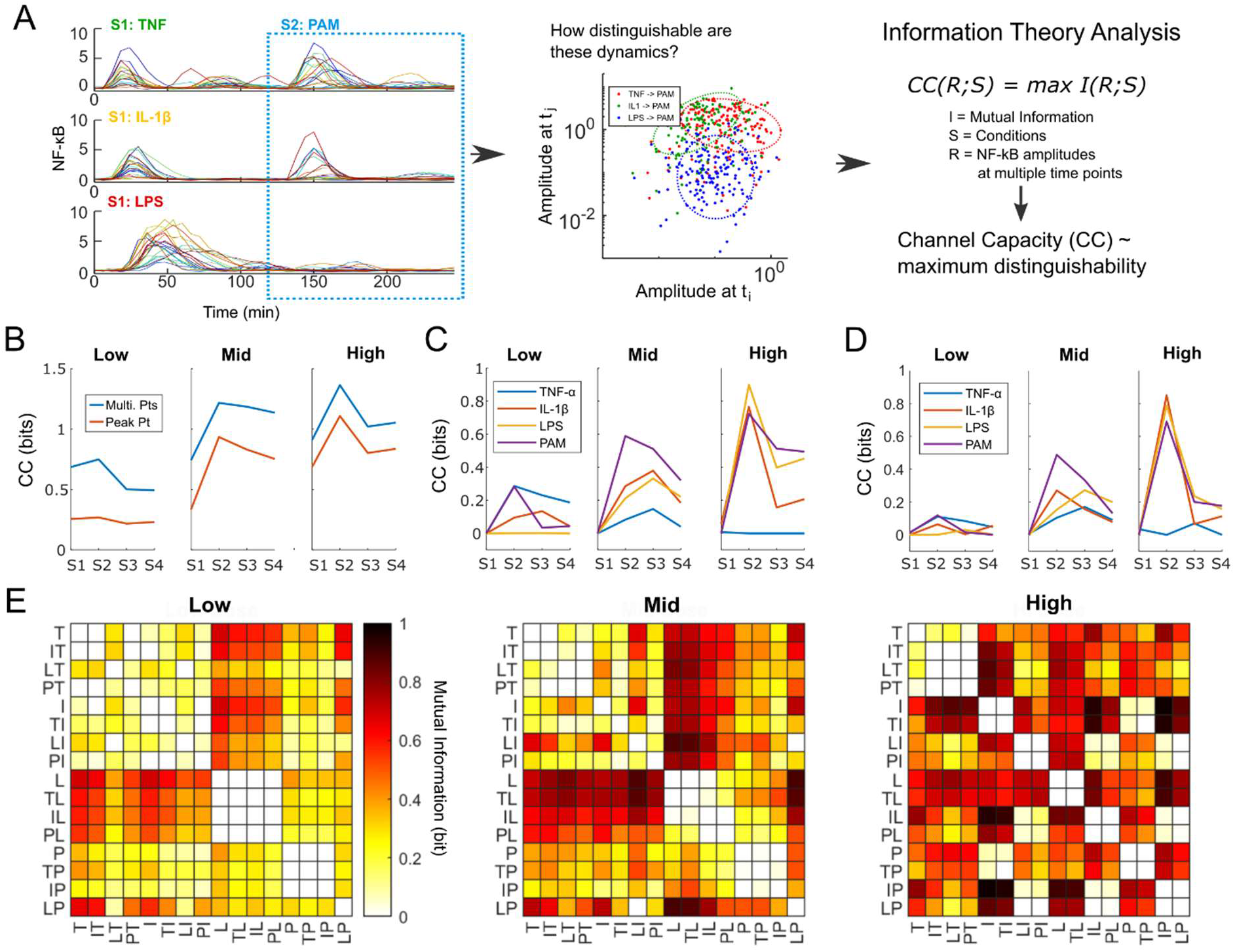
Information about prior stimulus history is reflected in the dynamics of subsequent NF-κB responses. A) Schematic representation of information theory analysis. Nuclear NF-κB levels at six different time points (20, 30, 40, 50, 70, and 90 min) from multiple conditions are used as inputs to calculate the mutual information between conditions. Channel capacity (CC) represents the maximum mutual information between conditions, which is comparable to the maximum distinguishability between condition. B) Distinguishability among all samples evaluated at each sequence order for each dose. CC is calculated from the 6-dimension vector (blue line) and compared to the CC from a single feature (red line) C) CC among all samples with the same ligand at each sequence position calculated using single cell NF-κB dynamics. CC in S2 – 4 indicates how accurately the NF-κB network reflects the prior history in the response to the indicated ligand. D) As in C), CC among all samples with the same ligand at each sequence interval but calculated using a single feature. E) Mutual information (MI) between ligand response dynamics (S1 and S2 only). T, I, L, P indicates the order of the stimulus. For example, T vs. IT indicates the MI between the response to TNF-α following IL-1β and the naïve TNF-α response. MI of 1 indicates complete distinguishability between two conditions.

To investigate how prior history affected the response for each ligand, we quantified the CC for each ligand at positions S1–4. We grouped the samples based on ligand and sequence position and calculated the CC among the samples within each group (Fig 3C). Ligands unaffected by prior history would produce identical responses and a CC of zero, while ligands for which prior history changes activation dynamics would see an increase in CC at S2–4. We found that the CC specific to each ligand generally rose at S2 and remained elevated at S3–4. In other words, more distinct response behaviors are present in S2–4, indicating that cell’s response to a specific ligand is significantly changed based on the cell’s prior history. However, TNF-α at high dose and LPS at low dose gained little information from prior history, which reflected our observations that prior history only weakly attenuated signaling in those samples (Fig 2A, C, Supp. Fig 5A, C). Nonetheless, the general trend of increased CC at S2–4 compared to S1 suggests that the NF-κB network encodes information about prior history in subsequent responses.

We also noted that the dynamics of the NF-κB response play a major role in accurate information transmission from prior history. When we compared the CC using response amplitudes at multiple timepoints to the CC using a single feature (the response amplitude when the mean was at its peak), we found that the CC from a single feature (Fig 3B red lines, 3D) are substantially lower than the CC from the dynamic measurement (Fig 3B blue lines, 3C). This indicates that alteration of NF-κB activation dynamics plays a role in transmitting information about prior history^31^.

We then investigated which ligand responses were most distinguishable from each other by calculating the mutual information between a pair of ligand responses. We focused on the naïve responses to a ligand at S1 and following another ligand at S2, resulting in a comparison of 16 conditions for each dose (Fig 3E). In the comparison matrix, mutual information patterns at low and mid dose were primarily driven by differences between TNF-α or IL-1β response dynamics and LPS or PAM response dynamics (e.g., comparing TNF-α and LPS). However, response to IL-1β or PAM following LPS (LI or LP) also were distinguishable from almost every other response at mid dose. At high dose, the pattern of mutual information changed such that all TNF-α responses became highly distinguishable from other samples. Likewise, the naïve and TNF-α exposed responses to IL-1β, LPS, and PAM also became distinguishable from same responses following either IL-1β, LPS, or PAM (e.g., TI vs LI or TL vs IL). This shift in mutual information patterns between low, mid, and high doses suggests that fundamentally distinct mechanisms could potentially mediate the effects of prior history in these dose ranges. Overall, these mutual information analyses confirmed that the NF-κB response is distinguished based on ligand sequence at the single cell level.

### Prior stimuli attenuate the subsequent NF-κB response in a ligand- and dose-dependent manner

To study how information about prior history is stored in the NF-κB network, we investigated how different stimuli produced different patterns of attenuation (Fig 1C). At all three dose ranges, TNF-α signaling was only weakly attenuated by prior stimulus, while the attenuation of LPS, PAM, and IL-1β signaling varied depending on the dose and identity of the prior ligand (Fig 2A-C, Supp. Fig 5). LPS, PAM, and IL-1β signaling all utilize a MyD88-dependent signal transduction pathway, including the shared signaling intermediary IRAK1 (Fig 1B)^10^. IRAK1 has been reported to regulate itself through autoinhibitory phosphorylation, which limits subsequent activation of IRAK1 by other stimuli^23^. Thus, we hypothesized that prior MyD88-dependent signaling attenuates subsequent signaling in the same pathway, but that TNF-α is independent from this inhibition.

To test this hypothesis, we focused on how a single prior ligand affects the following response, i.e., how the S1 ligand response changes the S2 ligand response (Fig 4A-C). We found that, following TNF-α stimulus, the response to MyD88-dependent ligands was weakly attenuated (Fig 4D, blue). Likewise, the response to TNF-α following MyD88-dependent stimuli was weakly attenuated (Fig 4D, green). In contrast, MyD88-dependent ligands attenuated subsequent signaling by other MyD88-dependent ligands in a dose-dependent manner (Fig 4D, red). At high and mid doses, exposure to MyD88-dependent ligands resulted in significantly attenuated signaling from other MyD88-dependent ligands compared to previous TNF-α exposure. Taken together, these results indicate that a prior history of TNF-α signaling minimally affected MyD88-dependent signaling and vice versa, while a prior history of MyD88-dependent signaling inhibited the response to other MyD88-dependent ligands in a dose-dependent manner.

**Figure 4:**
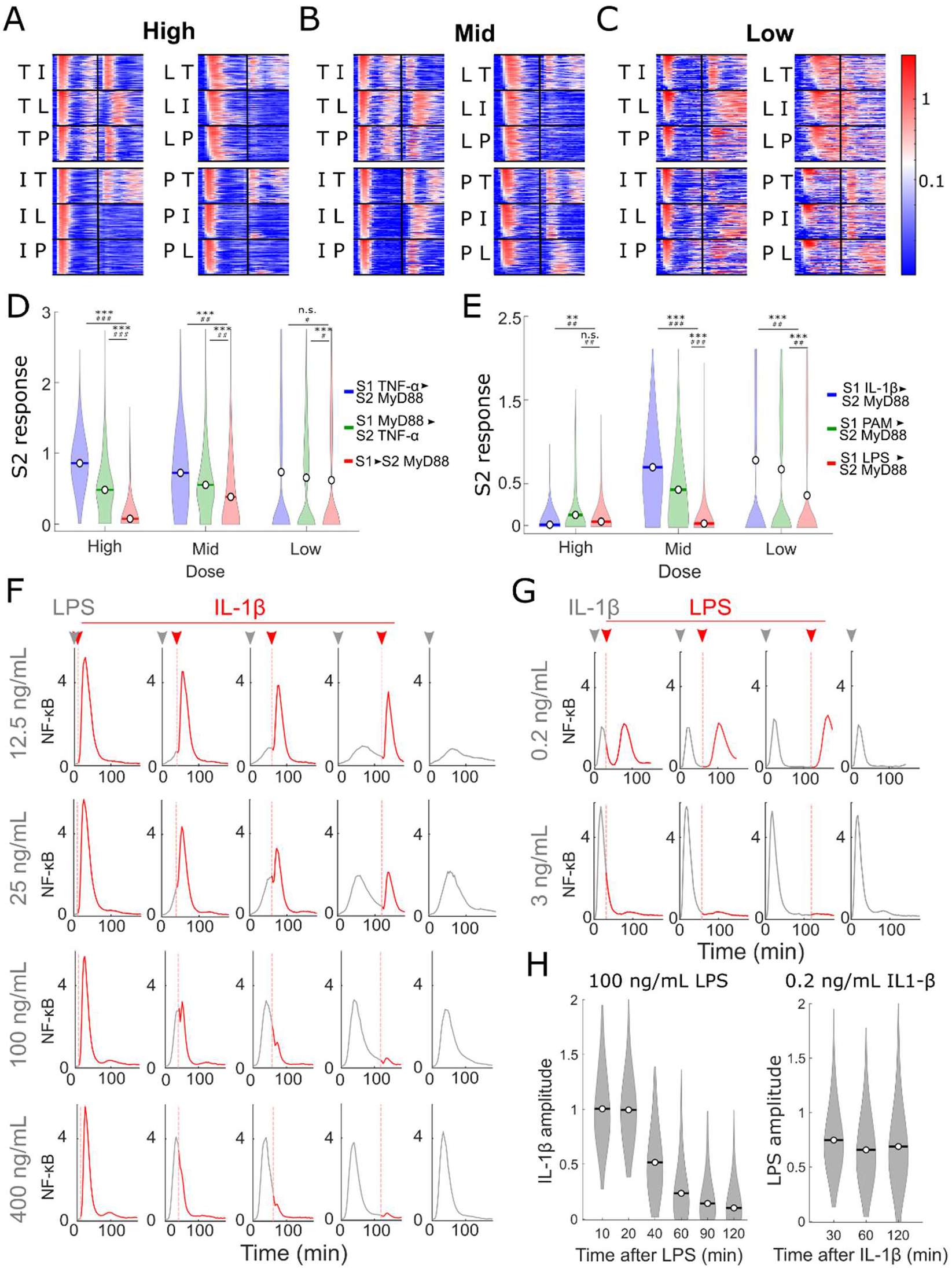
Ligand and dose specific effects of prior history differentiate TNF-α from MyD88 dependent ligands and differentiate among MyD88 dependent ligands. A-C) NF-κB response dynamics over 2 hours of stimulus for each ligand normalized to the mean amplitude of the naïve (S1) response. 50 single-cell traces randomly selected for each condition. All sequences of S1 and S2 ligands shown. All 9 sequences shown at high (A), mid (B), and low (C) dose. D) Violin plot comparing the normalized S2 responses of the MyD88-dependent ligands (LPS, PAM, IL-1β) following either TNF-α (blue) or another MyD88-dependent ligand (red) or the TNF-α response following a MyD88 dependent ligand (green) from > 650 cells per condition. Open circle and line show the mean. E) Violin plot comparing the normalized S2 MyD88-dep responses following IL-1β (blue), PAM (green), and LPS (red) stimulus at high, mid, and low doses from > 340 cells per condition. F) Plot of mean trace for conditions where LPS is provided at 0 min (gray arrowhead) and switched to 3 ng/mL (high dose) IL-1β after the indicated time (red arrowhead). Gray region of trace in each plot indicates NF-κB response during LPS stimulus interval and red region of trace indicates NF-κB response after replacement with IL-1β. Dashed red line indicates time of replacement. Plot at left of each row indicates the LPS response without switching to IL-1β (entirely gray trace). Each mean trace represents > 100 single cells. G) Plot of mean traces for conditions where 0.2 and 3 ng/mL (mid and high dose) IL-1β are provided at 0 min and switched to 100 and 400 ng/mL (mid and high dose) LPS, respectively, after the indicated time. Gray region of trace in each plot indicates NF-κB response during IL-1β stimulus interval and red region of trace indicates NF-κB response after replacement with LPS. Dashed red line indicates time of replacement. Plot at left of each row indicates the IL-1β response without switching to LPS (entirely gray trace). Each mean trace represents > 100 single cells. H) Violin plot comparing the normalized response for 3 ng/mL IL-1β following 100 ng/mL LPS or the response for 100 ng/mL LPS following 0.2 ng/mL IL-1β. Each plot is derived from > 100 cells per condition. Open circle and line show the mean. Bonferroni corrected Wilcoxon rank sum test p-value > 10^−2^ (n.s.), < 10^−2^ (*), < 10^−3^ (**), < 1*10^−4^ (***). Fold change difference between sample means > 1 (#), > 1.25 (##), or >4 (###).

If shared negative feedback is the primary cause of attenuation for subsequent MyD88-dependent signaling, each MyD88-dependent ligand should equally attenuate subsequent MyD88-dependent ligands. Although LPS is known to also utilize a MyD88-independent module mediated by TRIF and TRAM^35,36^, we found that the MyD88-independent pathway for LPS had minimal influence in these cells, as knocking out MyD88 was sufficient to abolish all response to LPS (Supp. Fig 7). Thus, we expected LPS, PAM, and IL-1β to equally inhibit the response to each other. At high dose, all three MyD88-dependent ligands indeed strongly attenuated subsequent responses (Fig 4E). In contrast, at mid and low doses, only LPS strongly attenuated subsequent MyD88-dependent signaling (Fig 4E, red), while IL-1β and PAM allowed significantly stronger subsequent responses (Fig 4E, blue, green). Thus, at high dose, attenuation between LPS, PAM, and IL-1β occurred symmetrically, while at mid and low doses, attenuation became asymmetric. Prior LPS stimulus inhibited subsequent IL-1β/PAM response but not vice versa. Similarly, when we compared the JNK responses to MyD88-dependent ligands following either IL-1β or LPS stimulus, we saw that symmetric attenuation took place at high dose, but at mid dose, only LPS maintained strong attenuation of MyD88-dependent JNK activation (Supp. Fig 8). These data reproduced the asymmetry in attenuation observed in our NF-κB measurements and suggest that asymmetric prior history effects may be broadly applicable in multiple inflammatory signaling pathways. Despite the highly shared pathways between LPS, PAM, and IL-1β, prior LPS effects differ from prior PAM or IL-1β effects in a dose dependent manner.

### Slow LPS-dependent negative feedback induces distinct attenuation and response behavior to the subsequent stimulus

While autoinhibition of IRAK1 can explain symmetric attenuation at high ligand dose^23^, as IRAK1 is shared by each of the MyD88-dependent ligands (Fig 1B), it could not explain our results at mid and low doses. Asymmetric cross-attenuation at mid and low doses suggests the existence of an additional negative feedback mechanism which would be more strongly activated by LPS stimulation than by IL-1β or PAM.

To study the characteristics of asymmetric attenuation of MyD88-dependent signaling, we examined how rapidly attenuation takes place upon stimulation with LPS. The timescale of attenuation can inform where in a signaling network the feedback acts. For example, rapid attenuation is unlikely to be driven by transcription and translation of downstream feedback genes. We stimulated cells with various doses of LPS (12.5 – 400 ng/ml), then stimulated the cells with high dose of IL-1β (3 ng/ml) after 10-120 minutes of LPS stimulus (Fig 4F, Supp. Fig 9A). Attenuation of IL-1β signaling by high dose LPS (400 ng/ml) was fast and strong, rapidly suppressing the subsequent IL-1β response at all times except the shortest time interval (10 min). As IRAK1 is shared in the early part of the signaling pathway, this observation was consistent with rapid autoinhibition of IRAK1. However, following lower doses of LPS, the IL-1β response became gradually attenuated depending on duration of LPS stimulus (Fig 4F, H).

On the other hand, when we stimulated first with IL-1β, then LPS, we did not observe gradual attenuation. Similar to high dose LPS, high dose IL-1β still produced immediate and strong attenuation of the LPS response, suggesting autoinhibition of IRAK1 still plays a major role in subsequent attenuation (Fig 4G, Supp. Fig 9B). Increasing duration of stimulus with mid dose IL-1β, however, had no impact on attenuation of LPS signaling (Fig 4G). To compare the difference between prior stimulation with LPS and IL-1β more clearly, we normalized the responses to the second stimulus to the corresponding naïve responses (Fig 4H). As expected, the response to IL-1β following LPS gradually decreased over time, while LPS response following IL-1β remained consistent over time. These results suggest that an additional activation-time dependent negative feedback process is differentially regulated by each MyD88-dependent ligand. This time dependence led us to hypothesize that this additional feedback response relies on NF-κB dependent gene expression.

### Ligand-specific attenuation in MyD88-dependent signaling depends on activation of IKK

To test whether NF-κB translocation and subsequent gene expression is necessary for asymmetric and ligand-dependent attenuation, we targeted the signaling intermediary IKK. IKK controls the activation and translocation of NF-κB into the nucleus through degrading the inhibitory protein IκBα (Fig 1B). Using PS1145, a reversible small molecule inhibitor of the IKK-β subunit^36,37^, we blocked signaling downstream of IKK activation. Due to the reduced activity of IKK, pretreating cells with 40 μM PS1145 significantly reduced NF-κB translocation by LPS stimulation (Supp. Fig 10). To test the impact of IKK inhibition for attenuation of subsequent signaling events, we washed cells to remove the drug after LPS stimulation and restimulated with 3 ng/mL (high dose) IL-1β. Cells treated with PS1145 showed significantly stronger NF-κB responses to subsequent IL-1β stimulus compared to untreated cells (Fig 5A). Thus, aspects of NF-κB signaling downstream of IKK activation, e.g., NF-κB nuclear translocation and NF-κB-mediated gene expression, play a major role in LPS-dependent attenuation of subsequent signaling. Through these inhibition studies, we show that asymmetric attenuation of MyD88-dependent signaling depends on IKK activation and subsequent NF-κB nuclear translocation, suggesting that this asymmetry depends on NF-κB-mediated gene expression.

**Figure 5:**
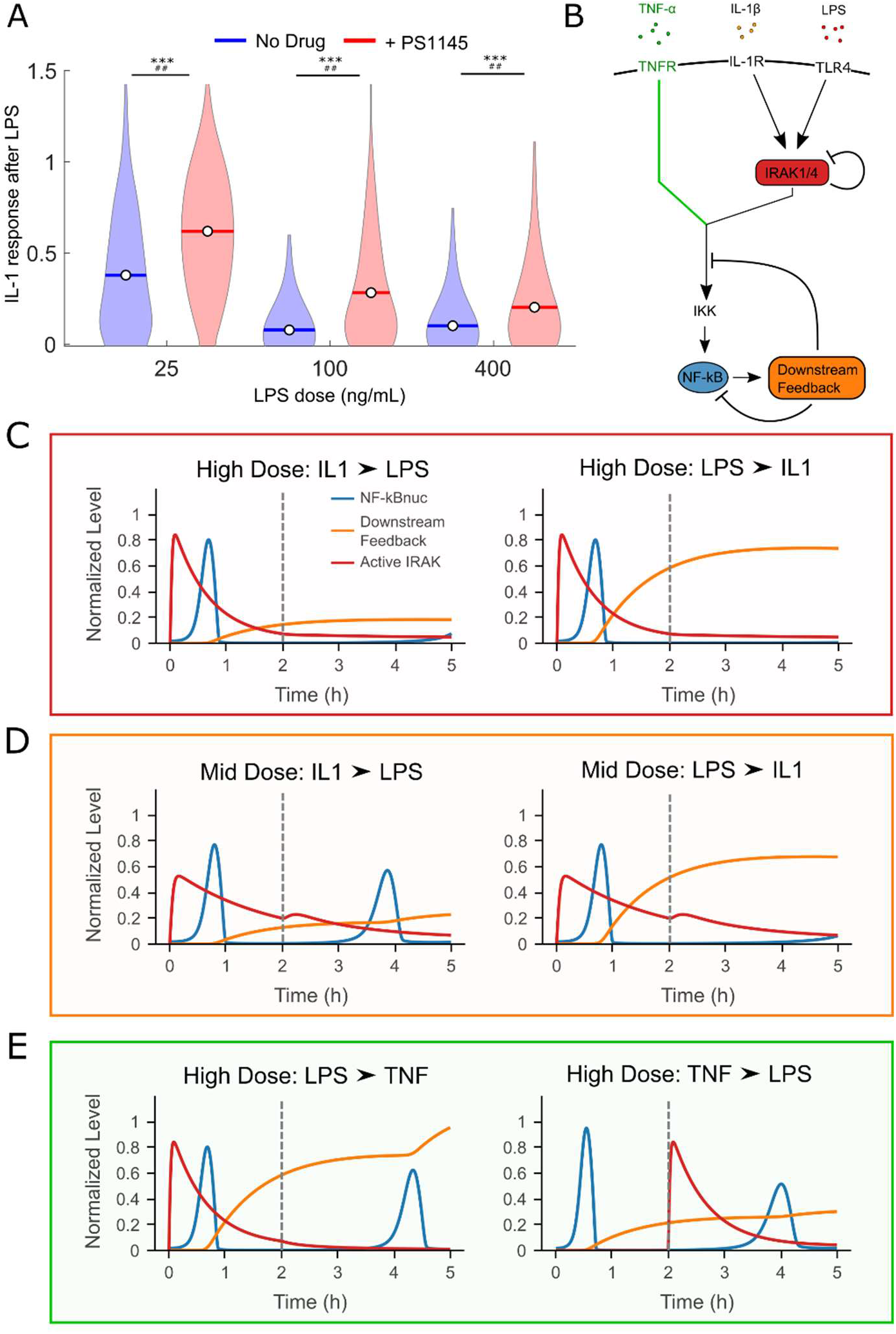
Differential regulation of downstream feedback controls ligand specificity of tolerance. A) Violin plot comparing IL-1β maximum response following LPS treatment normalized to naïve for untreated (blue) and PS1145 pre-treated (red) cells. Pre-treated cells were exposed to 40 μM PS1145 for 90 minutes prior to stimulus with LPS at the indicated concentration. Cells were fed twice with media after 4 hours of LPS treatment to remove PS1145 and 3 ng/mL IL-1β added. Each condition shown from > 120 single cells. B) Diagram illustrating the NF-κB network model used for the simulation. Two negative components, IRAK1 autoinhibition and nuclear NF-κB dependent attenuation, are highlighted in red and orange. The TNF-α signaling pathway (green) utilizes different kinases to activate IKK than the MyD88 dependent ligands. C—E) Simulated network responses to different sequences of stimuli. The blue lines show the dynamics of nuclear NF-κB, the red lines for active IRAK1, and the orange lines for the downstream feedback component. Gray dashed vertical line indicates time of similar replacement of first ligand with second (2 hours for each simulation).

### Mathematical modeling with two negative feedback motifs reproduces ligand and dose-specific tolerance effects

Our data give rise to a model where, at high dose, IRAK1 auto-inhibition results in symmetric attenuation of Myd88-dependent signaling, while at moderate and low doses, differential transcription of downstream negative regulators produces asymmetric attenuation. To study whether a network topology with these two motifs is sufficient to reproduce our observed prior history effects, we incorporated these two feedbacks into the NF-κB network model (Suppl. Information) and studied the change in network dynamics when stimulated with different ligand sequences.

To focus on the role of these two negative feedbacks, we minimized the network topology by converging all kinases not involved in negative feedback or the translocation of NF-κB^38^. Then, we expanded this minimal NF-κB model by adding network components connecting three receptors (TNFR, IL-1R, and TLR4) and incorporating autoinhibition of IRAK1 and ligand-dependent inhibition downstream of NF-κB (Fig 5B). Even with these expansions, our model uses only ~20 parameters and successfully reproduced our experimental observations (Fig 5C-E). At high dose of IL-1β or LPS, strong activation of IRAK1 resulted in rapid inactivation of itself, which prevented NF-κB activation by subsequent MyD88 ligands (Fig 5C). However, TNF-α was unaffected by this inactivation (Fig 5E).

In contrast, at mid dose, IL-1β induced weaker activation of IRAK1 and resulted in modest inactivation of itself, allowing activation of IRAK1 by subsequent LPS stimulus (Fig 5D). Partial inactivation of IRAK1 by LPS stimulus combined with induction of transcriptional feedback prevented subsequent MyD88-dependent signaling (Fig 5D). Thus, in this dose range, differential engagement of downstream feedback plays a critical role in differentiating LPS and IL-1β signaling and promoting asymmetric response. Additionally, we simulated other six combinations of sequential stimuli (Supp. Fig 11), which reproduced the remaining experimental results. Our simulation demonstrates how a simple network motif with a few negative feedbacks acting on different nodes can retain information about stimulus history and coordinate subsequent inflammatory signaling.

### MyD88-dependent ligands differentially regulate NF-κB response genes associated with negative feedback

Our computational and experimental results suggest that NF-κB-induced negative feedbacks are differentially regulated by MyD88-dependent ligands. To confirm this model, we profiled gene expression through RNA sequencing following 2 h of stimulation with mid dose LPS, PAM, or IL-1β. Compared to unstimulated cells, we found a total of 609 differentially expressed genes (DEGs) following LPS stimulus, 166 following PAM stimulus, and 108 following IL-1β stimulus (Table ST1). Almost all DEGs induced by IL-1β and PAM were also induced by LPS, while DEGs by IL-1β and PAM showed little overlap (Fig 6A). Differences in gene expression between these three ligands were primarily driven by magnitude of up or down-regulation, rather than regulation of different genes (Fig 6B). In general, upregulation of gene expression by LPS was stronger than upregulation by PAM, which was itself stronger than by IL-1β. These differences in the magnitude of gene expression suggest that MyD88 dependent ligands indeed differentially regulate expression of NF-κB response genes despite highly shared pathways.

**Figure 6:**
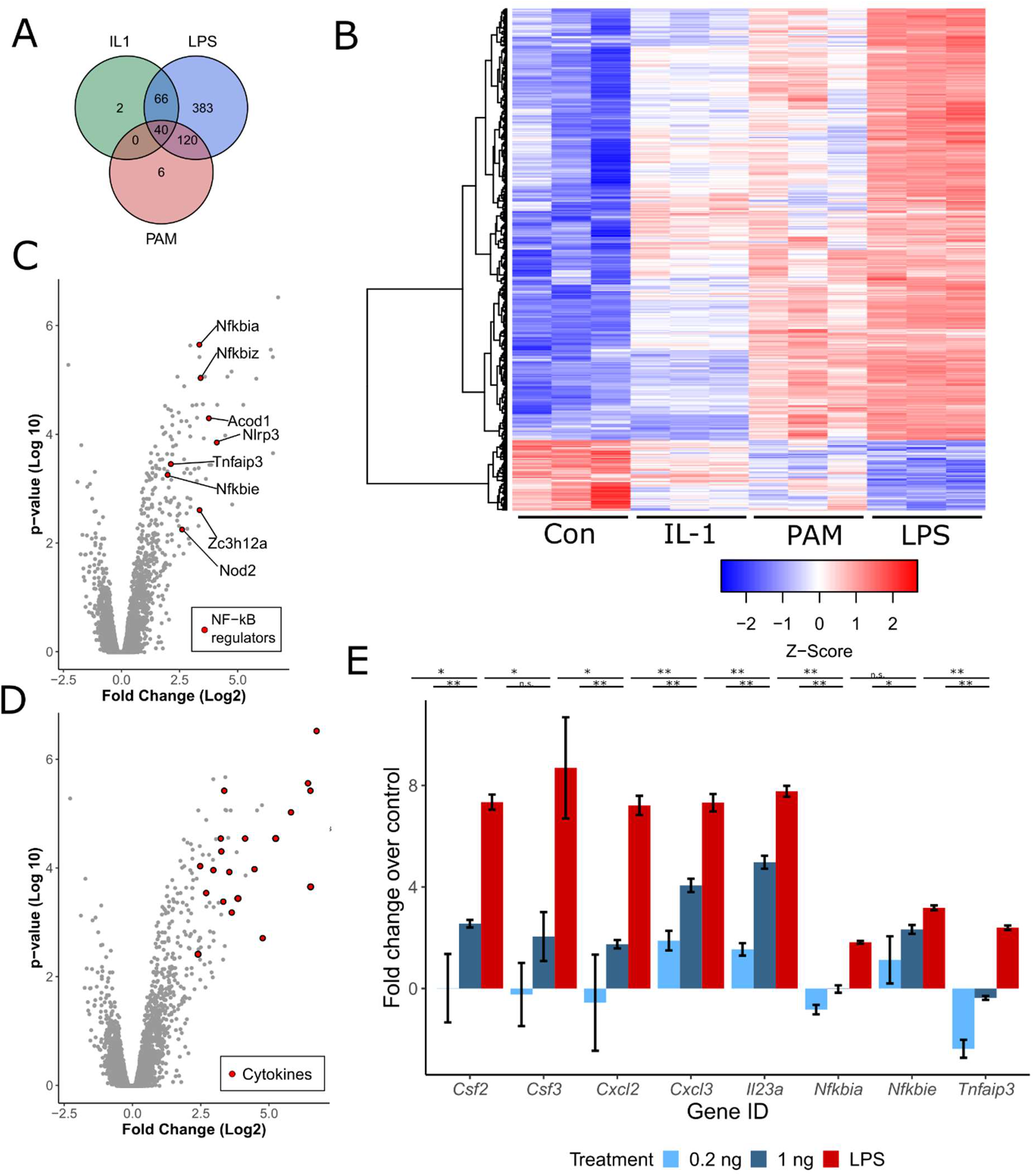
Myd88-depenent genes differentially regulate downstream cytokines and negative feedback regulators. A) Venn diagram showing overlap of differentially expressed genes (DEGs) between IL-1β, LPS, and PAM after 2 hours of stimulus. B) Heatmap of DEGs for MyD88-dependent ligand treated cells. RNA-sequencing was performed in triplicate. Each row shows the normalized expression (z-score) of a single gene. Dendrogram shows linkage based on Ward’s method. C-D) Volcano plot showing log2(fold change) and −log10(P value) for DEGs between LPS and IL-1β stimulus Among the DEGs with adjusted p-value < 0.01 and fold change > 4, genes annotated as NF-κB negative regulators (GO:0032088) (C) or genes annotated as cytokines (Gene Ontology, GO:0005125) (D) are colored in red. E) qRT-PCR data following for a subset of highly differentially expressed cytokines and NF-κB negative regulators stimulation with 0.2 ng/mL (light blue), 1 ng/mL (dark blue) IL-1β, or 100 ng/mL LPS (red). Gene expression is normalized to basal gene expression for unstimulated cells. Data shown as mean fold change over unstimulated cells +/− SEM from 3 replicates. Benjamini-Hochberg adjusted P value < 0.05 (*) or <0.01 (**).

We then focused on which genes were most differentially regulated by MyD88 ligands. We found that many known negative regulators of NF-κB signaling were upregulated 2- to 4-fold in response to LPS stimulus compared to IL-1β stimulus (Fig 6C). Many of these regulatory genes act to directly sequester NF-κB or inhibit the activities of shared upstream signaling components^39^. Thus, these negative regulators likely affect subsequent signaling by other MyD88 ligands. We also found that the most differentially expressed genes between LPS and IL-1β are signaling proteins, indicating that these transcriptional differences give rise to different functional outcomes between LPS and IL-1β signaling (Fig 6D). For example, some of the most differentially regulated genes were well-known proteins secreted by activated fibroblasts, including the growth factors *Csf2* and *Csf3* and the cytokines *Cxcl2* and *Cxcl3*^40–42^.

It is surprising that LPS and IL-1β induce different gene expression patterns despite similar intracellular pathways. Ligand-specific NF-κB activation dynamics may be involved in differentiating these expression patterns. LPS consistently produced a longer NF-κB activation duration than a comparable dose of IL-1β (Supp. Fig 12A). The duration of NF-κB activation has been shown to differentially regulate transcription of NF-κB response genes^15,16^, possibly explaining differences between IL-1β and LPS induced gene expression. However, longer activation duration also increases the total nuclear NF-κB over time.

To examine if total nuclear NF-κB, as measured by the area-under-the-curve (AUC) of the NF-κB response, can explain the differential gene expression by different ligands, we quantified gene expression in cells stimulated with mid dose IL-1β and LPS (0.2 ng/mL and 100 ng/mL, respectively) and a higher dose of IL-1β (1 ng/mL). Mid dose IL-1β produced lower AUC than mid dose LPS did, while 1 ng/mL IL-1β produced a similar AUC to mid dose LPS (Supp. Fig 12B). If higher total nuclear NF-κB explains stronger gene expression by mid dose LPS than by mid dose IL-1β, 1 ng/mL IL-1β would induce comparable downstream gene expression. Through reverse-transcription quantitative PCR (RT-qPCR), we profiled the transcription of three differentially expressed negative feedback regulators, *Nfkbia, Nfkbie,* and *Tnfaip3*. We found that for *Nfkbia* and *Tnfaip3*, expression was significantly increased in the 100 ng/ml LPS sample compared to both 0.2 ng/ml and 1 ng/ml IL-1β samples (Fig 6E). Thus, even with higher IL-1β concentration which induced comparable NF-κB response AUC to mid dose LPS, negative regulators of NF-κB are upregulated in LPS stimulation.

Similarly, we profiled five secreted proteins which were also highly differentially expressed between LPS and IL-1β stimulus, *Csf2, Csf3, Cxcl2, Cxcl3,* and *Il23a*. Each of these genes except *Csf3* was significantly upregulated following LPS stimulation compared to both IL-1β doses. These results suggest that AUC cannot explain the differential downstream expression we observed, but that NF-κB dynamic features, such as activation duration, drive the differential downstream expression between LPS and IL-1β. Overall, our downstream analyses demonstrate that each MyD88-dependent ligand differentially regulates downstream gene expression, and that differences in negative feedback expression can store prior ligand information to control subsequent NF-κB signaling.

## Discussion

Cells involved in innate immunity must interpret a complex and evolving milieu of extracellular cytokines and pathogenic signals. Despite the temporal features of these challenges, how prior history of inflammatory stimulus reshapes cellular responses to subsequent stimuli remains unclear. Here, we combined microfluidics and live-cell tracking of canonical NF-κB signaling to track the effects of complex stimulus patterns on inflammatory signaling over the course of hours.

Our results showed that different levels of overlap between ligand pathways and negative feedback modules encode information about prior history and shape response to subsequent ligands. In particular, TNF-α permitted signaling from subsequent ligands and is least affected by prior stimulus history. In contrast, prior history between the MyD88-dependent ligands is differentiated by dose-dependent engagement of shared IRAK1 autoinhibition and ligand-dependent production of downstream negative feedback proteins. The combination of these three network features is sufficient to differentiate subsequent ligand responses following a prior history of ligands with highly shared pathways, like IL-1β and LPS, based on activation dynamics to subsequent stimuli.

Thus, we propose a model of acute “memory” of prior history in the NF-κB network where ligand-specific engagement of negative feedbacks remodels nodes of the NF-κB network shared with subsequent ligands (Fig 7). This memory of prior history reshapes the response to the subsequent stimulus, resulting in significantly different NF-kB translocation dynamics. While these activation dynamics have been extensively shown to control transcriptional outcomes and cell fate^15,16,43^, we show that these dynamics can reflect state changes due to prior stimuli as well., In future study of innate immune memory, the role of network remodeling and signaling dynamics should therefore be considered as potential regulatory mechanisms for biological function.

**Figure 7:**
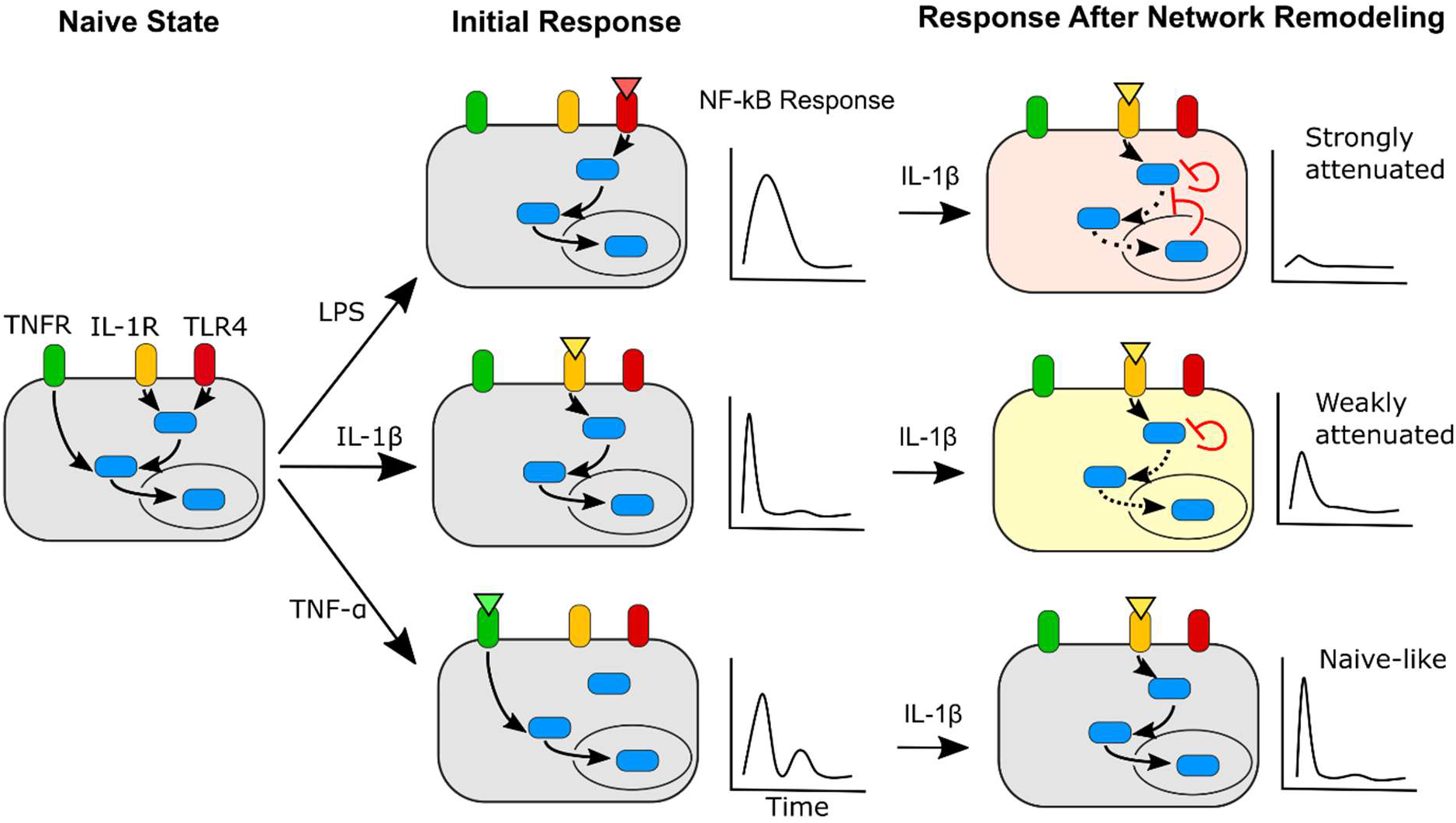
Exposure to inflammatory ligands remodels the NF-κB network to alter subsequent ligand responses. Naïve cell (gray) activates in response to different inflammatory ligands, which each remodel the NF-κB network in a characteristic manner. LPS (red cell) induces upstream and downstream negative feedback, IL-1β (yellow cell) primarily induces upstream negative feedback, and TNF-α induces negative feedback which primarily acts orthogonally to the other ligands. As a result, response to a subsequent IL-1β stimulus becomes attenuated in a ligand specific manner and produces “memory”-informed NF-κB responses.

Our finding that acute prior history effects are encoded in the dynamic NF-κB response also presents a novel framing of innate immune memory. We report that, in addition to epigenetically mediated effects on the timescale of days, innate immune memory can also be encoded by NF-κB dynamics regulated through rapid remodeling of the upstream signaling network. In myeloid cells with cell type-specific negative feedback modules like IRAK-M or the MyD88 splice variant MyD88S^44,45^, the regulation of innate immune memory by feedback-dependent alteration of NF-κB dynamics may be further enhanced. The regulation of NF-κB dynamics at longer timescales and in contexts relevant to other immune phenotypes like trained immunity or priming remains an open question that needs to be addressed.

## Acknowledgements

We thank laboratory members for critical reading and discussion of this work. We also thank Markus Covert (Stanford) and Sergi Regot (Johns Hopkins) for providing the cell line used in this study, and Alexander Hoffmann (UCLA) and Roy Wollman (UCLA) for providing the information theory code that our analyses were based on. AGW is supported by the NIH MSTP training grant T32GM07281. This work is supported by NIH grants R01GM128042, 75N93019C00041 and Army Research Office grant 73231-EL (ST).

## Contributions

AGW and MS designed and performed the microfluidics experiments. AGW and MS analyzed the microfluidic data with help from NT. AGW performed the gene expression analyses with help from MS. MS built the mathematical models and information theory analyses. ST supervised the work. All authors contributed to writing of the manuscript.

## Materials and Methods

### Cell culture

RelA^-/-^ NIH3T3 immortalized mouse embryonic fibroblasts (3T3s) stably expressing RelA-DsRed, JNK-kinase translocation reporter-mCerulean3 (JNK-KTR)^46^, and histone 2B-green fluorescent protein (H2B-GFP) were cultured with Dulbecco’s Modified Eagle Medium – High Glucose (DMEM; Gibco) supplemented with 10% fetal bovine serum (Omega Scientific), 1% GlutaMAX (Gibco), and 100 u/mL penicillin-streptomycin (Gibco) in tissue-culture treated flasks. Cells were cultured in a tissue culture incubator maintained at 37°C and 5% CO_2_. Cells were passed prior to reaching 100% confluency and maintained for no more than 15 passages. On the day of the experiment, cells were harvested with trypsin, washed with complete medium, and resuspended at ~5*10^6^ cells/mL in FluoroBrite DMEM (Gibco) with the same supplements to reduce background fluorescence.

### Microfluidic device design and fabrication

A previously designed and published cell culture device was utilized for automated cell culture and ligand stimulus^24^. The design contains 14 unique stimulus inputs and 64 independently controlled cell culture chambers measuring 3.5 × 0.8 × 0.035 mm, where each can load more than 500 cells. Master molds for this chip were fabricated by patterning photoresist deposited on silicon wafers through multilayer soft lithography^47^. Microfluidic devices were fabricated by pouring polydimethylsiloxane (PDMS; Momentive, RTV-615) on the control and flow master molds and bonding these two layers. Control layer wafers were poured with 66 g PDMS (10:1 monomer to catalyst), air bubbles removed under vacuum, and cured at 80°C overnight to make a ~2 cm thick PDM slab with the control pattern grooved on the bottom. Flow layer wafers were poured with 15 g PDMS (10:1 monomer to catalyst) and spun at 2200 rpm to achieve a thickness of ~50 μm and cured at 80°C for at least 1 hour. After curing, holes intended for control pins were punched in the control layer, both PDM layers were treated with oxygen plasma (Harrick, PDC-001), aligned using a custom stereomicroscope, and the aligned chip were baked at 80°C overnight. After bonding, holes intended for fluid input and output were punched; then the chip was bonded to a glass slide through plasma treatment and baking. A detailed fabrication protocol can be found in our previous publications^24,27^.

### Microfluidic experiment setup

Device control layer inputs were connected to pneumatic solenoid valves with electronic controller boxes. By actuating different sets of valves, flow pathways in the microfluidic device can be directed from a particular input to a particular chamber using pre-written Matlab scripts and a custom-developed graphic user interface (GUI). The device was mounted on a microscope (Nikon) and cell chambers were filled with 0.25 mg/mL fibronectin (Millipore) in sterile pH 7.4 phosphate buffered saline (PBS, Gibco), and incubated overnight at room temperature. Subsequently, chambers and channels were flushed with complete medium to replace the fibronectin, then the temperature, humidity, and CO_2_ in the live imaging apparatus (Life Imaging Services) were set to 37°C, 100% humidity, and 5% CO_2_ to optimize cell culturing in the microfluidic device. Cells were loaded at approximately 50% confluency to optimize tracking efficiency, and cells were allowed to settle and equilibrate for 5 hours prior to start of stimulation and imaging.

### Stimulus conditions

Four ligands, mouse tumor necrosis factor alpha (TNF-α; R&D Systems, aa 80-235), mouse interleukin 1 beta (IL-1β; R&D Systems, 401ML010CF), ultrapure lipopolysaccharide (LPS) from E. coli (InvivoGen, tlrl-3pelps), and PAM2CSK4 (PAM; InvivoGen, tlrl-pm2s) were utilized in this study. Based on experimental quantification of NF-κB translocation following titration of each ligand, we selected high, mid, and low doses of each ligand with comparable activation (TNF-α: 90, 30, 3 ng/mL; IL-1β: 3, 0.2, 0.05 ng/mL; LPS: 400, 100, 12.5 ng/mL; PAM: 1, 0.1, 0.01 ng/mL). For each set of high, middle, and low dose ligands, all non-repeating combinations of the four ligands were supplied at 2-hour intervals, producing 24 conditions per dose over 8 hours. One condition was maintained as a positive control (mid dose TNF-α, IL-1β, LPs, PAM) and one condition maintained as a negative control (4 feedings of complete media). For other experimental conditions, ligands were provided and switched at the indicated dose at the indicated time. Ligand dilutions were made from stock solutions stored at −80°C immediately prior to stimulus, stored on ice during the duration of the experiment, and delivered to the chip through polyetheretherketone tubing (VICI, TPK.505). Input pressure was maintained at 4 psi to prevent shear stress on cells during feeding. For IKK inhibition experiments, PS1145 (Tocris, 4569) was diluted in complete media to 40 μM. Cells were pretreated with PS1145 for 90 minutes, then exposed to media containing PS1145 and LPS for 4 hours, washed for 30 minutes in complete media, and stimulated with IL-1β (3 ng/mL). Other detailed protocol for the microfluidic experiment can be found in our previous publications^27^.

### Image acquisition and analysis

Epifluorescence images were acquired using a Nikon Ti2 microscope enclosed within a temperature-controlled incubator (Life Imaging Services). Images were captured at 20X magnification through a complementary metal-oxide semiconductor camera (Hamamatsu, ORCA-Flash4.0 V2) every 6 minutes. Each chamber position was imaged for p65-DsRed (555-nm excitation, 0.5-1 s exposure time), H2B-GFP (485-nm, 50-100 ms), and/or KTR-JNK-mCerulean3 (440-nm, 100 ms). No photobleaching or phototoxicity was observed over the course of the imaging process. For the time resolved experiments switching from LPS to IL-1β, imaging was conducted every 3 minutes instead in order to increase the temporal resolution of the trace.

Prior to image processing, background fluorescence and dark frame images were taken for flat field correction. Nuclear and cytoplasmic DsRed and/or mCerulean3 fluorescence for single cells were evaluated over the course of the experiment by analyzing time course fluorescence images with custom developed software (MATLAB). Briefly, H2B-GFP images were used to segment the nuclear region for each cell, whose positions were tracked over the entire sequence of time course images. Combining these single cell trajectories with the DsRed and mCerulean3 images, we quantified the median nuclear fluorescence in the nucleus, which represented the nuclear NF-κB level, and normalized this fluorescence to the median cytoplasmic fluorescence evaluated from a ring of cytoplasm located around the segmented nuclear image^48^. To quantify the background fluorescence, a few small regions without cells were randomly selected, and their mean fluorescence were evaluated and subtracted from the corresponding fluorescence measurement. The resulting traces were processed using another custom-developed analysis software to remove traces displaying cell death, division, or other features which impact data quality. Only traces which were complete over the entire course of each experiment were retained for subsequent analysis.

Key trace features were extracted using custom software (MATLAB). The frame of the maximum RelA or JNK-KTR response in a stimulus interval was identified using a trace smoothed with the lowess method with a span size of 3 to reduce noise from cell movement, slight changes in imaging focus, or background fluctuations. Frames identified from the smoothed trace were then used to identify the true maximum fluorescence in the un-smoothed trace. To account for the possibility of oscillations in nuclear translocation, multiple local maxima were allowed with a minimum distance between maxima of 5 frames (30 minutes). To distinguish true maxima from noise due to frame-by-frame fluctuation in nuclear fluorescence, we set the 95^th^ percentile of maxima identified from unstimulated cells as the cutoff and set all stimulus maxima below that cutoff to be zero. Area under the curve (AUC) for each stimulus interval was calculated by taking the trapezoidal approximate of the integral for each trace in the defined time interval.

### CRISPR-Cas9 knockout of MyD88

A *Myd88*-targeting guide RNA (5’-TCGCGCTTAACGTGGGAGTG-3’) was cloned into the pX330 plasmid backbone (Addgene Plasmid #42230) and transfected using electroporation (Lonza) into 3T3s. 48 hours post-transfection, single cells were sorted into a 96-well plate and allowed to grow into clonal populations. Screening by Sanger sequencing identified three clones with frameshift mutations in one or both copies of the gene. Successful knockout was confirmed with western blot probing for MyD88 (1° rabbit anti-MyD88 1:1000, Cell Signaling Technologies. 2° goat anti-rabbit DyLight 800 1:25000), following which the blot was stripped and reprobed for β-actin as a loading control (mouse anti-β-Actin Alexa Fluor 680, 1:1000). Blots were imaged on a LICOR scanner on the 700 and 800 nm channels.

### Cell retrieval from microfluidic device for downstream gene measurements

To facilitate retrieval of cells, the corner of the microfluidic device with the outlet was cut to expose the outlet channel. At the indicated time following stimulation, cells in the target chamber were treated with TrypLE Express (Gibco) for ~ 1 min to detach them from the treated surface, then sent to the outlet channel by washing with PBS. Detached cells accumulated at the outlet channel, were removed in a ~2 uL droplet by manual pipetting, and deposited in 10 uL ice-cold lysis buffer containing 0.1% Triton-X 100 and RNase inhibitor (Takara) and stored at −80°C until further processing. Approximately 1500 cells were retrieved per replicate per condition.

### Library preparation and RNA-sequencing

Sample prep for RNA-sequencing followed the SMART-Seq2 pipeline for single cells. Briefly, cell lysate was incubated at 72 °C with oligo-dT_30_VN to anneal, followed by the rest of the SMART-Seq2 reverse transcriptase mix and incubated at 42C for 90 minutes followed by 10 cycles between 50°C and 42°C to unfold secondary structure. Template switching using a modified TSO oligo (5′-AAGCAGTGGTATCAACGCAGAGTGAATrGrGrG −3′) provided a PCR handle on the 3’ end of the newly synthesized cDNA strand. 6 cycles of preamplification with KAPA HiFi (Roche), and purification with Ampure XP beads (1:1 ratio, Beckman Coulter) produced a purified cDNA library. Library prep was performed by the University of Chicago Genomics Facility using the Nextera XT procedure. Samples were then single end sequenced in the same facility on an Illumina HiSEQ4000 with a read length of 50 bp. Adapter trimming and read mapping to the reference mouse genome (GRCm38) was done using STAR using default parameters. Transcript abundance was quantified using featureCounts. Raw counts were normalized and differential gene expression identified using the R packages edgeR and limma. Differential genes were identified between IL-1β and untreated, PAM and untreated, LPS and untreated, and IL-1β and LPS using cutoffs of Benjamini-Hochberg false discovery rate (FDR) < 0.01 and log fold change > 1.

### cDNA synthesis and qPCR

Targeted reverse transcription and preamplification were done using a CellDirect One-Step RT-qPCR kit (Themo Fisher) as previously described. qPCR was performed with custom primer/probe sets (*Tnfaip3*), predesigned IDT PrimeTime probe assays (*Csf2, Csf3, Cxcl2, Cxcl3, Il23a, Gapdh*), or predesigned TaqMan probe assays (*Nfkbia, Nfkbie*). Ct values were calculated using software defaults and normalized to glyceraldehyde-3-phosphate dehydrogenase (GAPDH) expression to produce ΔCt values. ΔCt values were subtracted from the ΔCt values from control samples to calculate the ΔΔCt as a proxy for fold change expression over control.

### NF-κB Network Simulation

#### Building system of equations

To investigate if the two negative feedback model (Fig. 4C) is sufficient for the ligand history effect, we built a simplified network simulation. We extended the previous minimal NF-κB model^38^, which comprises three coupled differential equations each describing the dynamics of nuclear NF-κB (Eq. 1), mRNA of IκBα (Eq. 2), and cytoplasmic IκBα (Eq. 3). Previous study reports that nuclear NF-κB activates the transcription of downstream gene in a sigmoidal fashion with sharp threshold ^49,50^. Thus, we adapted the Hill function to describe mRNA transcription and applied a Hill coefficient of four to accurately describe the dynamics (Eq. 2). Our study involved various ligand stimuli, where each corresponds to different receptor and involves various cytoplasmic kinases for NF-κB activation. However, all signaling pathways converge on an essential mediator, IKK, prior to NF-κB translocation^11^. Upon activation from upstream stimuli, the neutral IKK becomes active, and degrades IκBα initiating the NF-κB translocation. The active IKK gradually turns into an inactive form, which then goes back to the neutral state over time^51^. We added two differential equations to describe this cycling of IKK (Eq. 10 and 11). Then, we incorporated the two negative feedbacks discussed in our study. Upstream of IKK, MyD88-dependent ligands (LPS or IL-1β) converge on another common kinase, IRAK1/4, which was shown to have auto-inhibitory negative feedback function reliant on aggregation^23^. The strong activation of IRAK1/4 facilitates its own inactivation, which eventually inhibits all MyD88 dependent stimuli after initial NF-κB activation. To integrate this important upstream negative feedback, we added IRAK1/4 activation and inactivation dynamics for each MyD88 dependent receptor (Eq. 6 — 9). To minimize variables, we assumed that the activation and inactivation rates by different receptors are same, and thus that IRAK1 kinetics depend only on the amount of each receptor in active state matters. Since the inactivation rate varied by the amount of active IRAK, we made the inactivation term non-linear, where the inactivation rate is proportional to the squared concentration of active IRAK. Another important negative feedback originates downstream of NF-κB. Other than IκBα, previous works report many downstream genes, which inhibit nuclear NF-κB in various ways^39^. Among them, several inhibitors target upstream of IKK, where many negative feedbacks including A20, SOCS-1/3, and Trim30α, repress the receptor activity and thereby hinder the activation of IKK. Hence, we added the expression of the downstream negative inhibitor (Eq. 4 and 5) and adjusted the IKK activation term in Eq. 10 to incorporate this effect. The Hill coefficient of 3 in this inhibition term includes the high cooperativity that may arise in the complex associations between multiple molecules in the upstream. For example, for A20 to be fully active, it not only needs to be dimerized but also needs other adaptor proteins to inhibit the phosphorylation of IKK^26^. Additionally, IKK has multiple phosphorylation sites, which may require multiple inhibitor complexes to successfully repress the IKK activation^52^. These complexities would likely contribute to the high cooperativity in the reactions happening in the upstream signaling cascade. Lastly, for the amount of activate ligand receptors, we normalized the dose range of ligand such that similar dose would activate similar number or ratio of receptors. For simplicity, we applied fast equilibrium approximation for the receptor dynamics, *i.e.*, at any given time the activity of receptor simply corresponds to the dose of ligand (Eq. 12 — 14). All receptors investigated in our study require multimerization to be active ^10,12^; hence, we used non-linear relationship between the dose and the active receptor. The system of equations for our model is listed below:

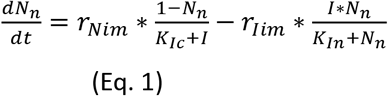

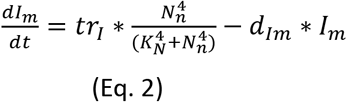

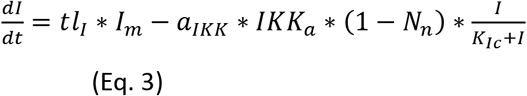

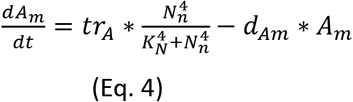

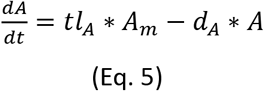

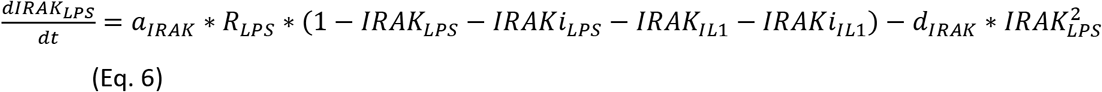

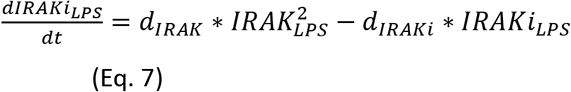

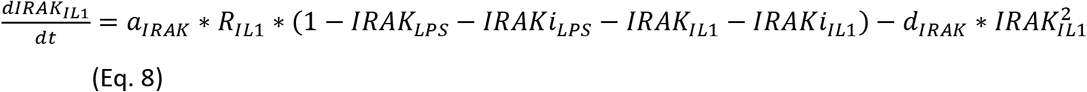

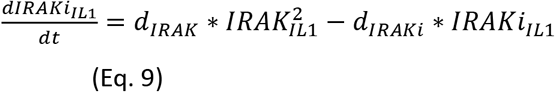

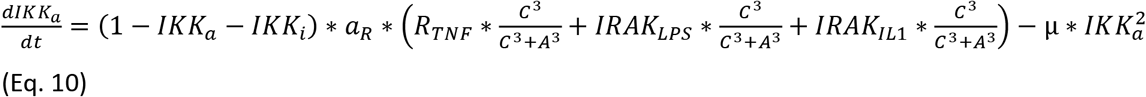

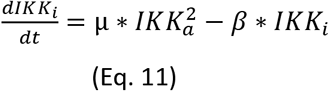

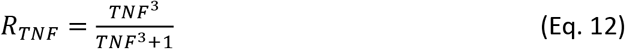

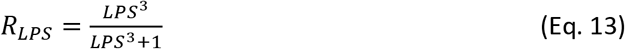

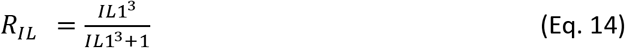

#### Values for parameters

Even though our model consists of the two negative feedbacks and multiple receptors, we managed to reduce the number of parameters to twenty. Roughly half of these are related to NF-κB and IκBα dynamics. The other half describes the newly added mechanisms, which involve dynamics of IKK cycling and negative feedback regulations. Since our model is based on the minimal model from the previous publications, we adapted parameters from them to where applicable. For the newly added components, we assumed or fitted the parameters to the period of NF-κB oscillation (~ 2h). The list of parameters and their values are described in the table ^24,38,53,54^.

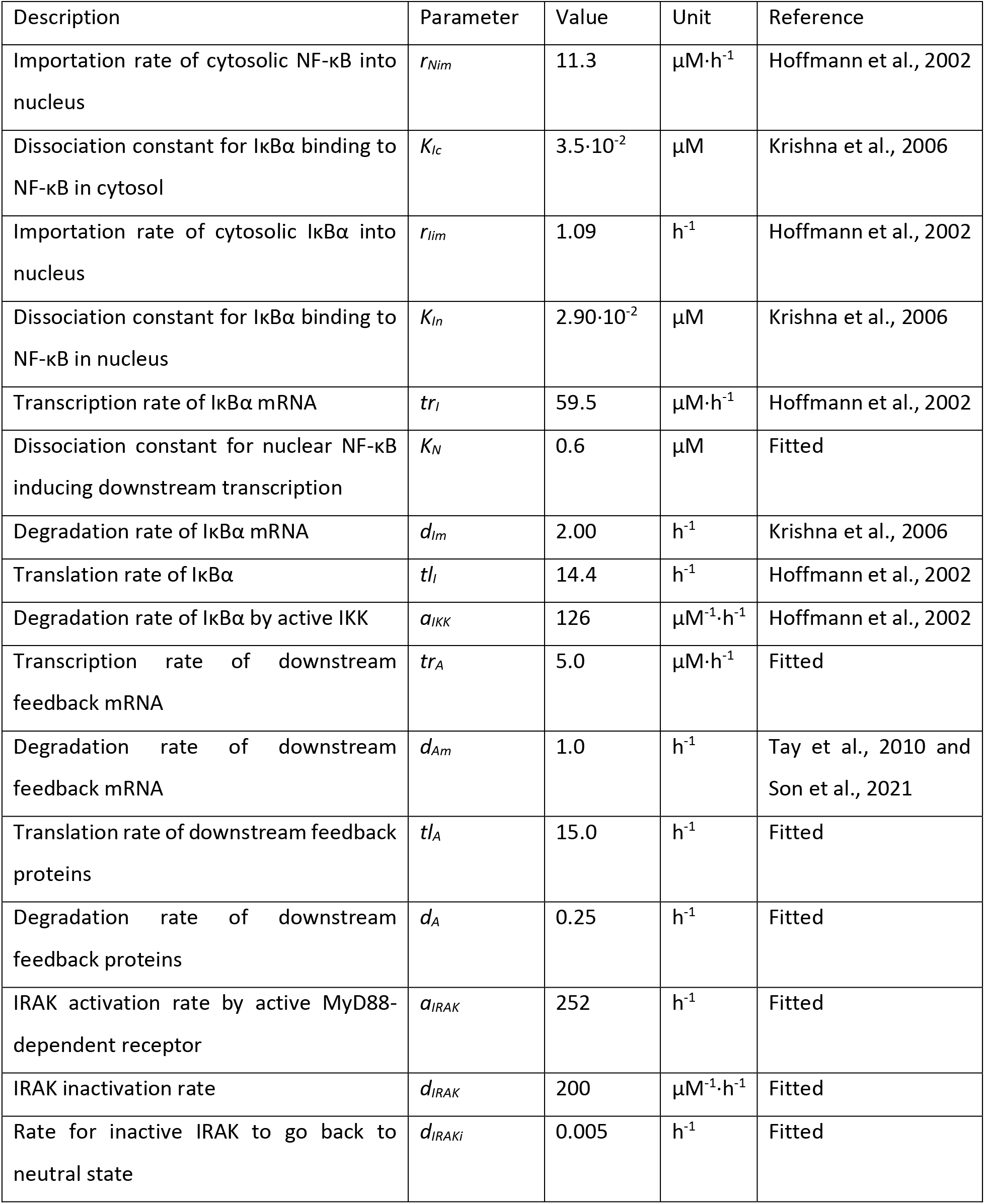
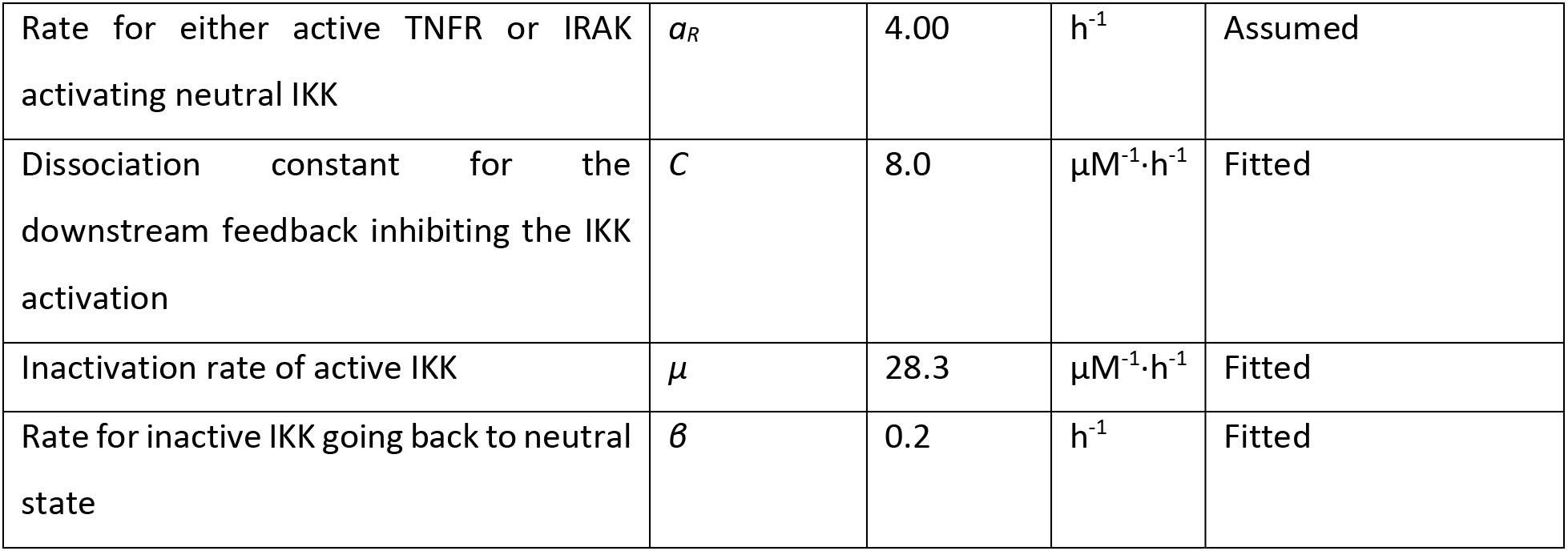

#### Running simulations

Computer simulations were performed using Python. The differential equations were integrated using *odeint* from *scipy.integrate* solver. To determine the basal stationary state of the network prior to stimulation, a short pulse of TNF-α was introduced at the beginning, then the dynamic of each component in the network was monitored up to 48 h after the pulse. After confirming the dynamics of all components became stationary, we stimulated the network with one of the first ligands (TNF-α, IL-1β, or LPS), then replaced it with another ligand after 2 h. Simulated dynamics of different components were plotted using Bokeh visualization library.

To simulate the difference in the expression level of the downstream negative feedback, we adjusted the dissociation constant for the inhibition of IKK activation (parameter *C*). If we added the different downstream expression parameters for each ligand, it would dramatically increase the number of parameters necessary to describe the dynamics of downstream negative feedback. Since all we needed was having different IKK inhibition strength from each ligand, we achieved the same effect by simply adjusting the dissociation constant for IKK inhibition. More specifically, for LPS stimulation, the dissociation constant was reduced by half, meaning the threshold for negative feedback molecules to inhibit the IKK activation is reduced by half. This way we could still monitor the effect from the different downstream negative feedback strength, while minimizing the number of parameters.

### Information Theory Analysis

For the information theory analysis, we employed the method and codes developed by Selimkhanov *et al.* ^31^. After obtaining the dynamic of NF-κB translocation in each cell, the nuclear NF-κB level at multiple time points were extracted and used as response (variable *R*) to evaluate the mutual information (variable *I*). Briefly, the mutual information is equal to the difference between the entropy of entire response (i.e., non-conditional entropy) from all samples and the sum of entropies from response in each sample (conditional entropy)^55^:

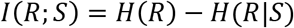

where *I* indicates the mutual or transfer information and *H* indicates the entropy. Hence, it indicates the reduction of uncertainty in ‘guessing’ which sample the response came from after observing the response. However, each sample may have different probability of happening. For example, in the case of this study, cells may be exposed a particular ligand sequence more frequently than other sequences. The conditional entropy can fluctuate depending the probability of each sample (or ligand sequence). However, it is still possible to evaluate what would be the maximum information transfer possible through the given system or NF-κB network. This is defined as channel capacity, *C*, and can be evaluated by finding a set of probabilities that would maximize the mutual information:

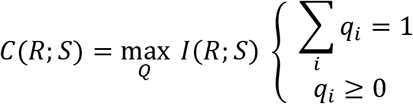

where *C* indicates the channel capacity, *Q* is a set of probabilities for *m* samples, [*q*_*1*_, *q*_*2*_,... *q*_*m*_]. Further details about the calculating entropies and how the mutual information was maximized can be found in the previous publication^31^. In this study, the NF-κB levels at multiple time points during each ligand interval in each sample were used as input (variable *R*) to calculate the channel capacity of NF-κB network in distinguishing a particular ligand at each step (S1-4) or prior history of ligand.

