## Supplemental figures for "NF-κB memory coordinates transcriptional responses to dynamic inflammatory stimuli"

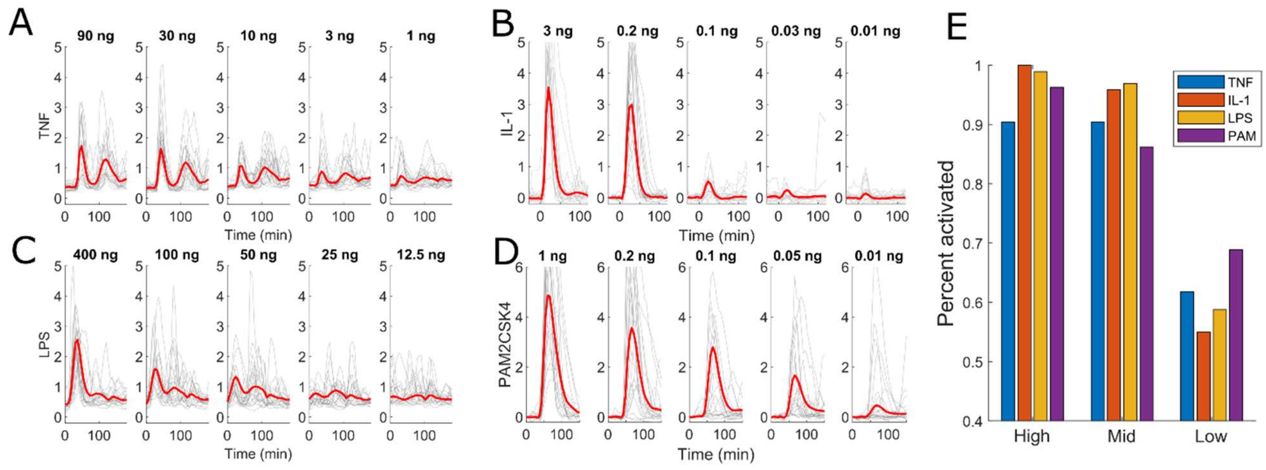


**Supplemental Figure 1:** NF-κB response to varying doses of the indicated ligands. 20 randomly selected single cells (gray lines) and mean (red line) are shown for each stimulus condition: A) TNF-α, B) IL-1β, C) LPS, or D) PAM. E) Percent activated for selected dose ranges – high (> 95% activated, TNF-α 90 ng/mL, IL-1β 3 ng/mL, LPS 400 ng/mL, PAM 1 ng/mL), mid (85-95% activated, TNF-α 30 ng/mL, IL-1β 0.2 ng/mL, LPS 100 ng/mL, PAM 0.2 ng/mL), low (50-70% activated, TNF-α 3 ng/mL, IL-1β 0.05 ng/mL, LPS 12.5 ng/mL, PAM 0.01 ng/mL). Maximal activation percentage for TNF-α is ~90% so high dose of TNF-α showed lower percent activation than high dose of other ligands.

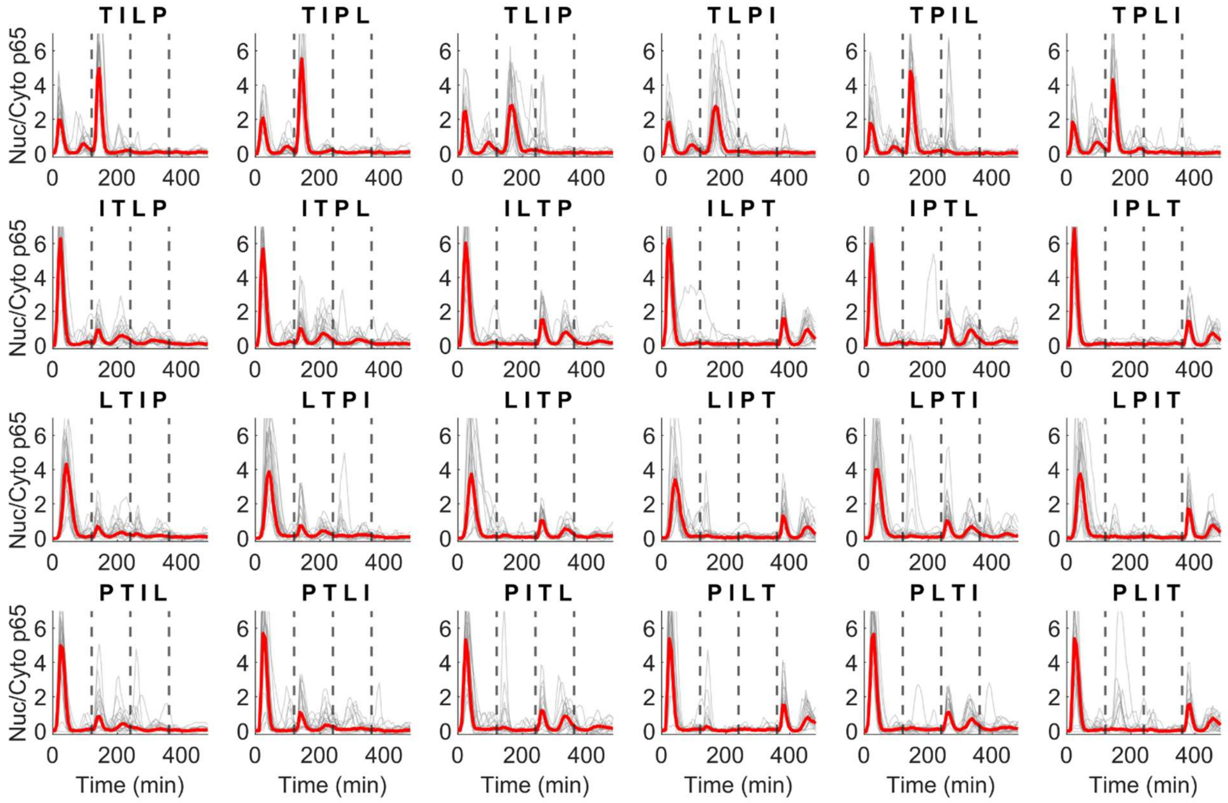


**Supplemental Figure 2:** NF-κB response to all 24 high dose combinations of TNF-α (T, 90 ng/mL), IL-1β (I, 3 ng/mL), LPS (L, 400 ng/mL), PAM (P, 1 ng/mL). 20 randomly selected single cells from > 200 cells (gray lines) and mean (red line) shown for each stimulus condition. Dotted lines indicate the timing for second, third, or fourth stimulus.

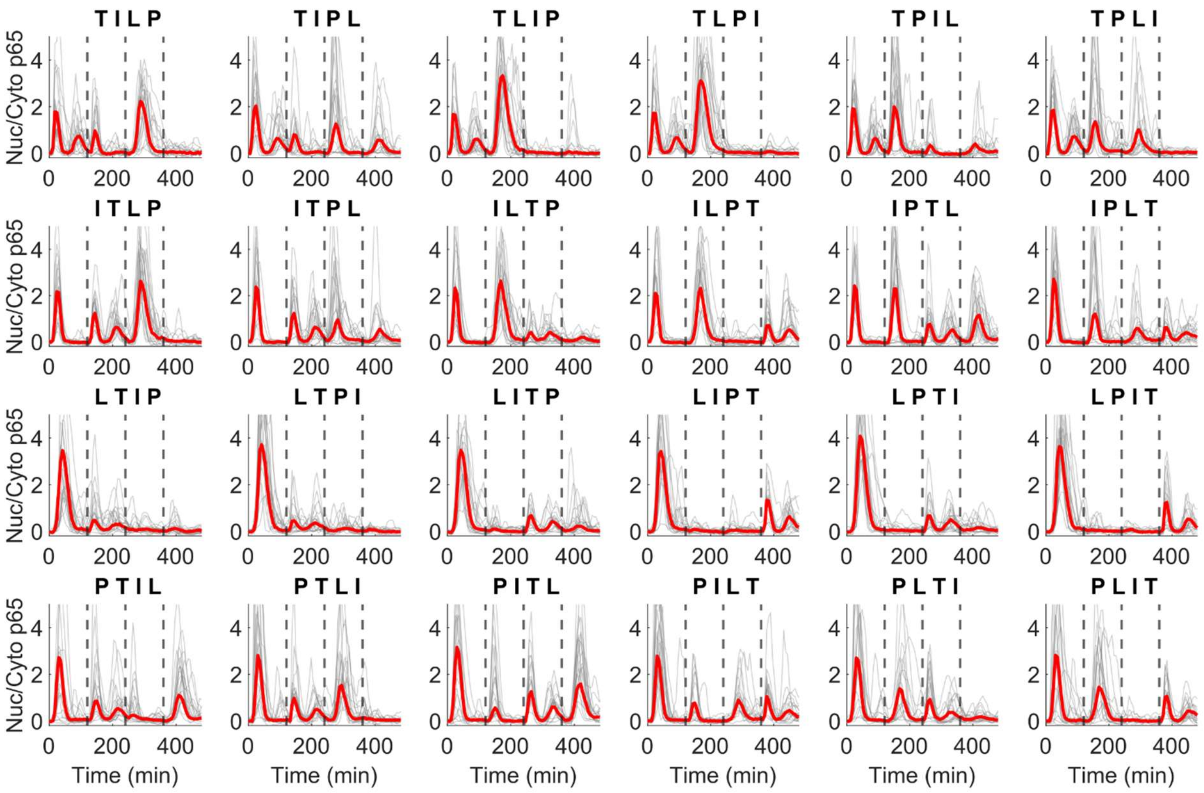


**Supplemental Figure 3:** NF-κB response to all 24 mid dose combinations of TNF-α (T, 30 ng/mL), IL-1β (I, 0.2 ng/mL), LPS (L, 100 ng/mL), PAM (P, 0.2 ng/mL). 20 randomly selected single cells from > 200 cells (gray lines) and mean (red line) shown for each stimulus condition. Dotted lines indicate the timing for second, third, or fourth stimulus.

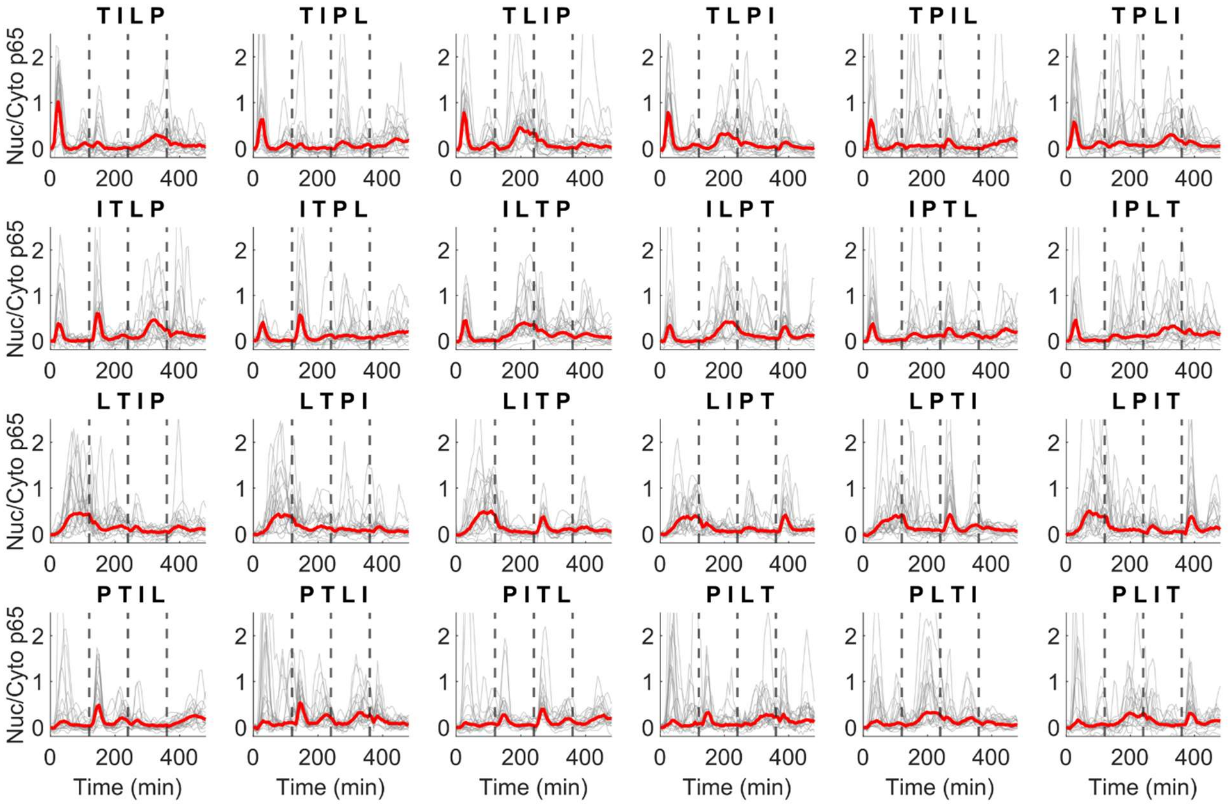


**Supplemental Figure 4:** NF-κB response to all 24 low dose combinations of TNF-α (T, 3 ng/mL), IL-1β (I 0.05 ng/mL), LPS (L 12.5 ng/mL), PAM (P, 0.01 ng/mL). 20 randomly selected single cells from > 200 cells (gray lines) and mean (red line) shown for each stimulus condition. Dotted lines indicate the timing for second, third, or fourth stimulus.

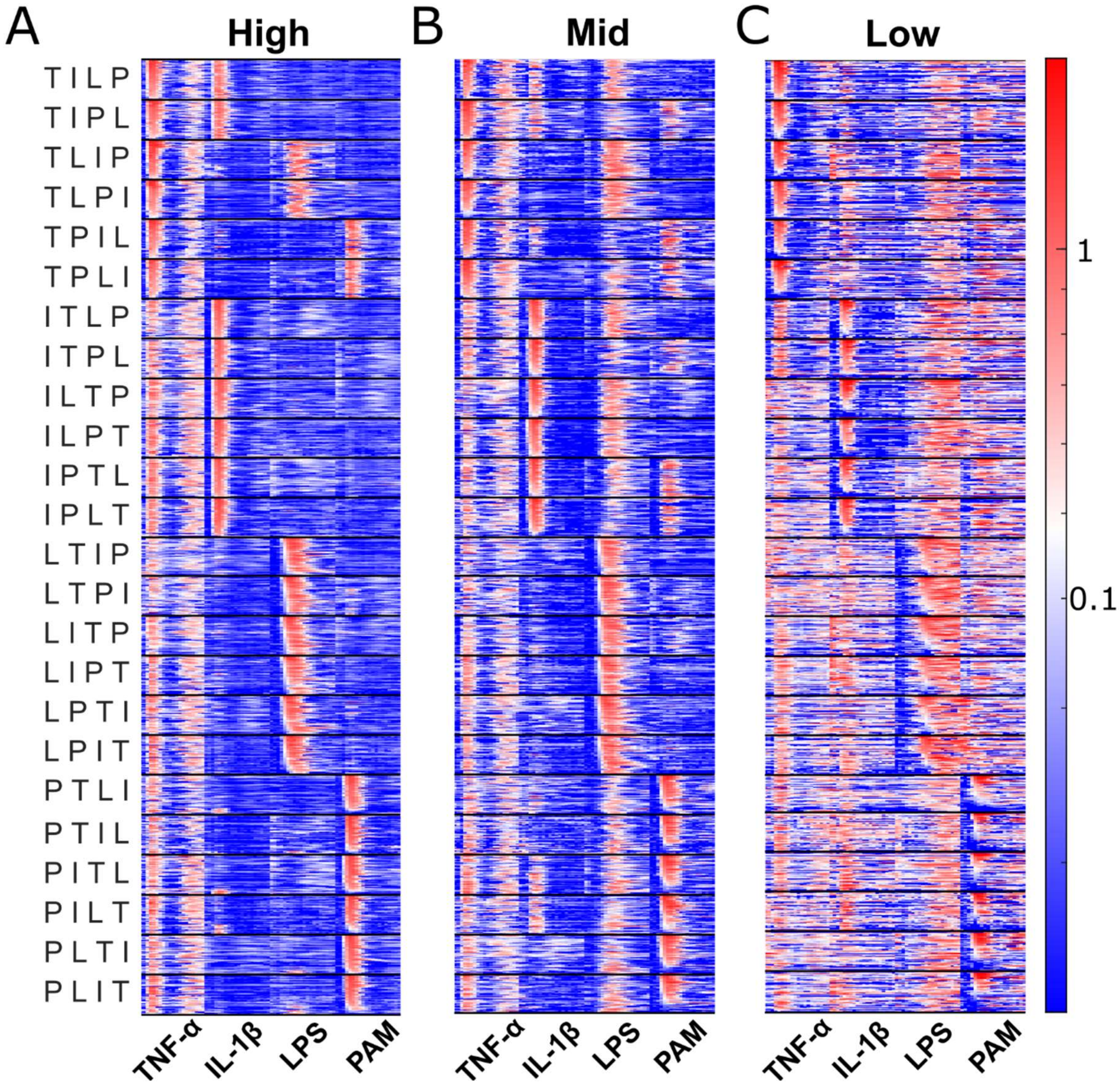


**Supplemental Figure 5:** NF-κB response dynamics over 2 hours of stimulus for each ligand normalized to the mean peak amplitude of the naïve (S1) response and plotted in heatmaps (50 randomly selected traces for each condition). Heatmap columns are arranged based on the stimulus ligand regardless of its order in the sequence. Stimulus orders are shown to the left of the first heatmap from S1 to S4, where T stands for TNF-α, I for IL-1β, L for LPS, and P for PAM. Plots shown for high (A), mid (B), and low (C) dose.

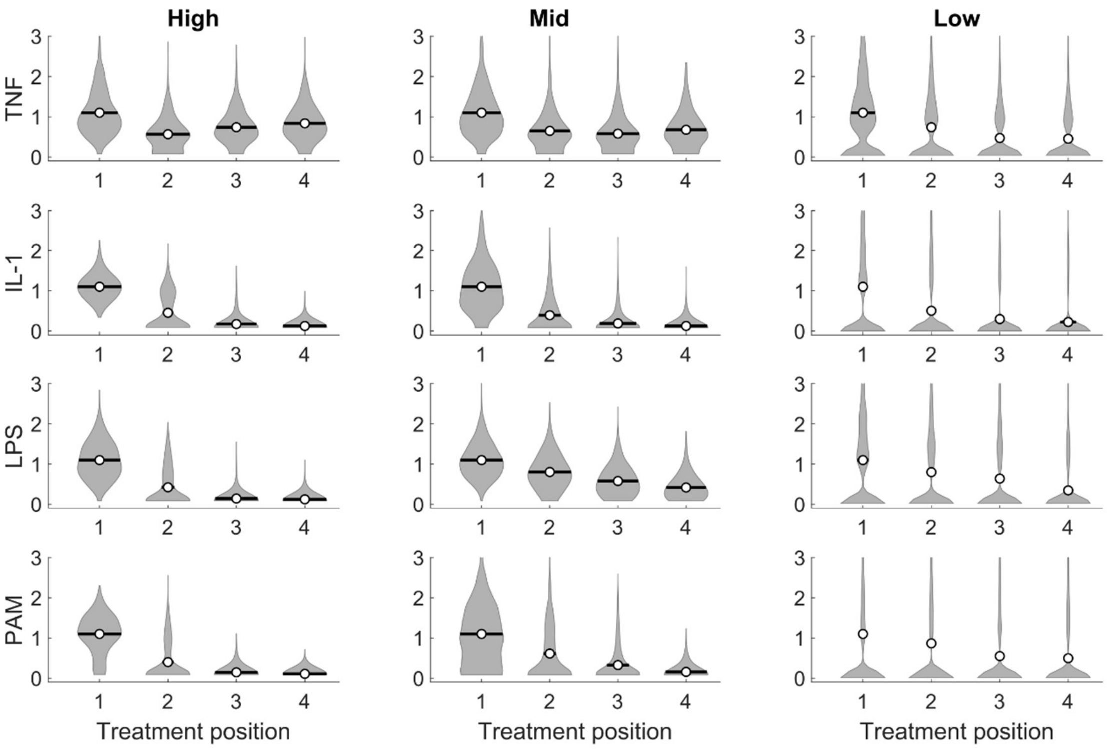


**Supplemental Figure 6:** Normalized peak responses to three different doses (low, mid, and high) of each ligand at different stimulus sequence (S1-S4). Responses in each time interval are normalized to the peak amplitude of the mean trace for the same ligand at S1. Each violin plot displays the responses of >500 single cells.

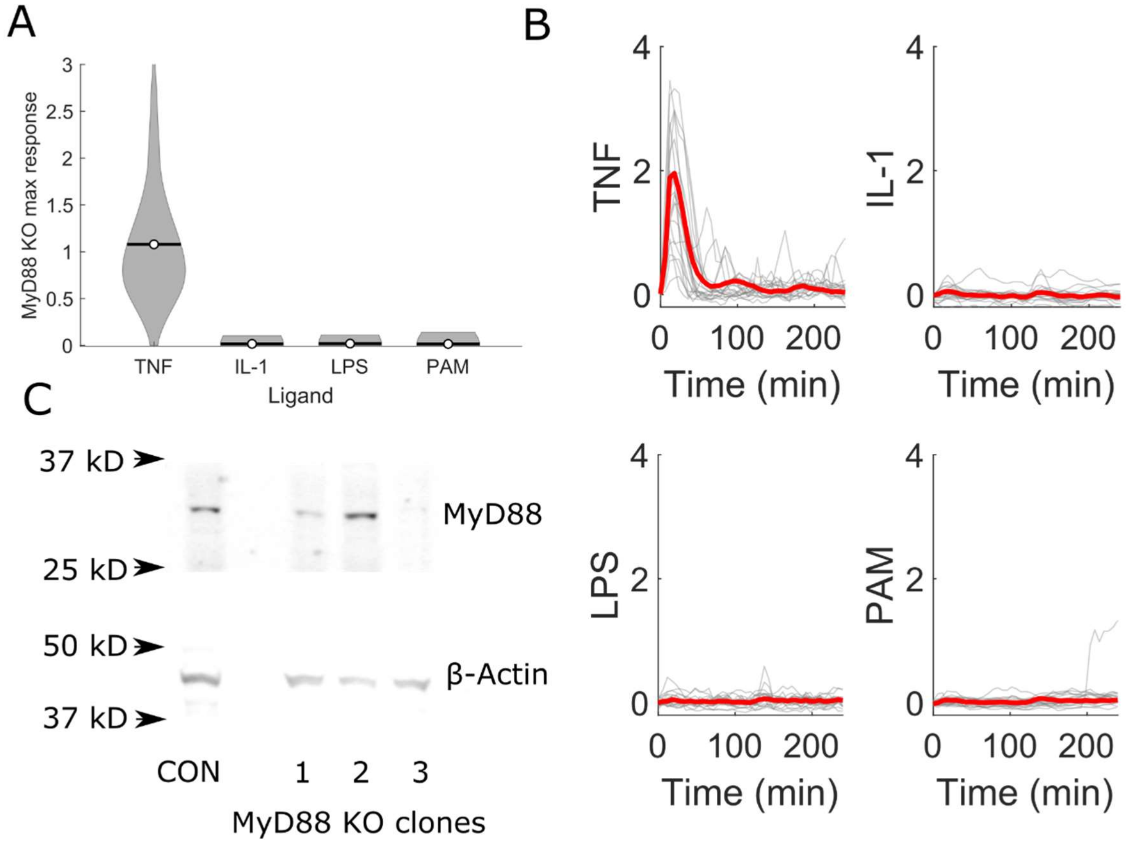


**Supplemental Figure 7:** MyD88 is critical for IL-1β, LPS, and PAM responses. A) Max response following stimulation by the indicated ligand in $MyD88^{-/-}$ cells normalized to max response following stimulation in WT cells. B) NF-κB traces for stimuli response shown in A). 20 randomly selected single cells from > 200 cells (gray lines) and mean (red line) shown for each stimulus condition. C). Western blot confirming successful knockout of MyD88 by CRISPR-Cas9. Three isolated single cell clones following CRISPR-Cas9 targeting of *Myd88* showing either frameshift mutations in both copies (clone 1, 3) or in one copy (clone 2) were probed for MyD88 and β-actin by western blot. Based on expression levels, MyD88 KO clone #3 was used for subsequent experiments. Full blots for both MyD88 and β-Actin can be found in supplemental figure 13.

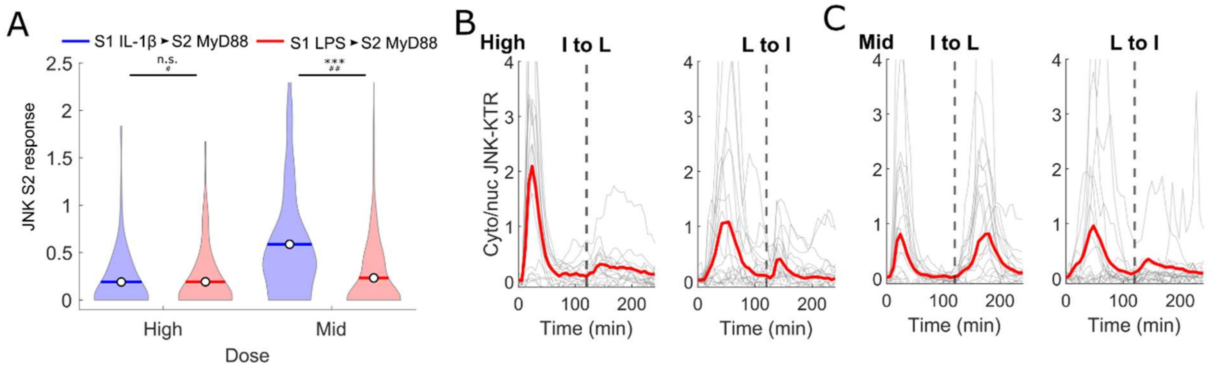


**Supplemental figure 8:** A) Violin plot comparing JNK response to MyD88-dependent ligands following 2 hours of IL-1β (blue) or LPS (red) stimulus at high and mid dose normalized to naïve response. Each condition shown from > 300 single cells. B) JNK response traces following stimulus with high dose LPS (L) or IL-1β (I) for 120 minutes switching to the other ligand (dashed vertical line). Data shown for 20 randomly selected single cells (gray lines) and population mean (red line) from > 200 cells. C) Same as B) at mid dose.

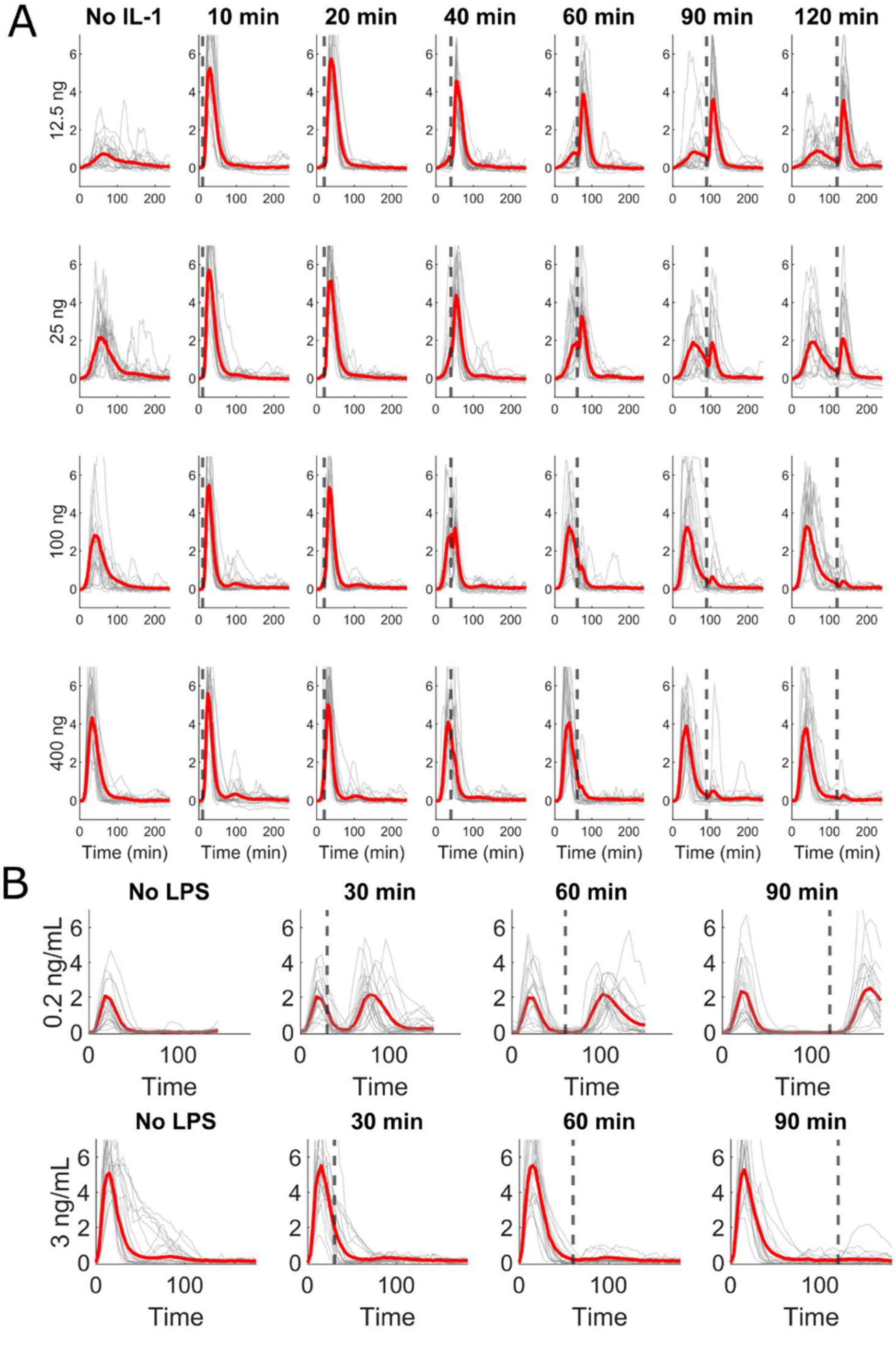


**Supplemental Figure 9:** NF-κB traces for time-resolved stimulation with A) LPS (various doses as indicated in the y-label) switching to IL-1β (3 ng/mL) or B) IL-1β (0.2 ng/mL or 3 ng/mL) switching to LPS (100 ng/mL or 400 ng/mL respectively). Dashed gray line indicates when the second stimulus was introduced. 20 randomly selected single cells (gray lines) and mean from > 100 cells (red line) shown for each stimulus condition.

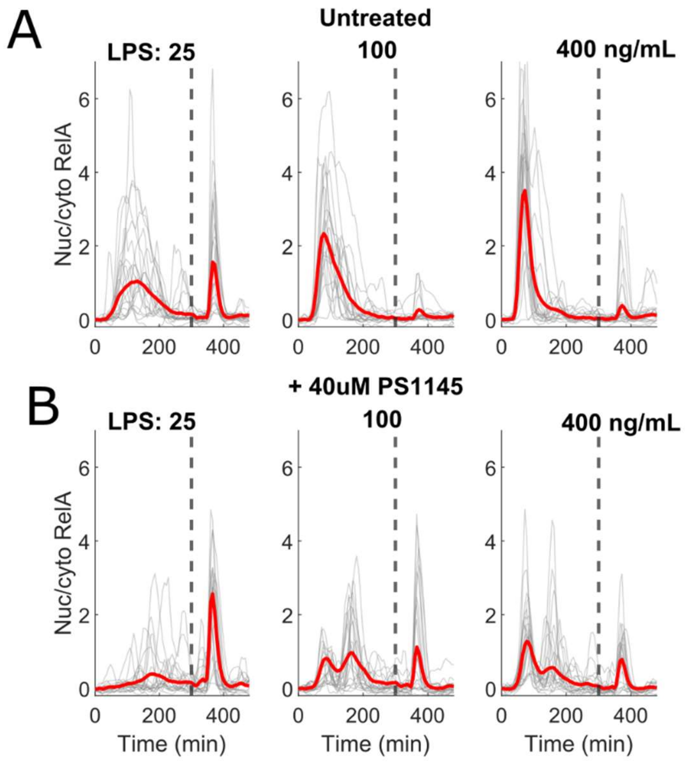


**Supplemental Figure 10:** A-B). NF-κB response traces from untreated cells (A) or cells pretreated with 90 minutes of 40 μM PS1145 stimulation (B). Cells were then exposed to LPS at the indicated dose for 240 minutes (+ 40 μM PS1145 for treated cells), washed with media twice for 15 minutes each, and stimulated with 3 ng/mL IL-1β. The dashed vertical line shows when IL-1β was supplied. Data shown for 20 randomly selected single cells (gray lines) and population mean (red line) from > 120 single cells.

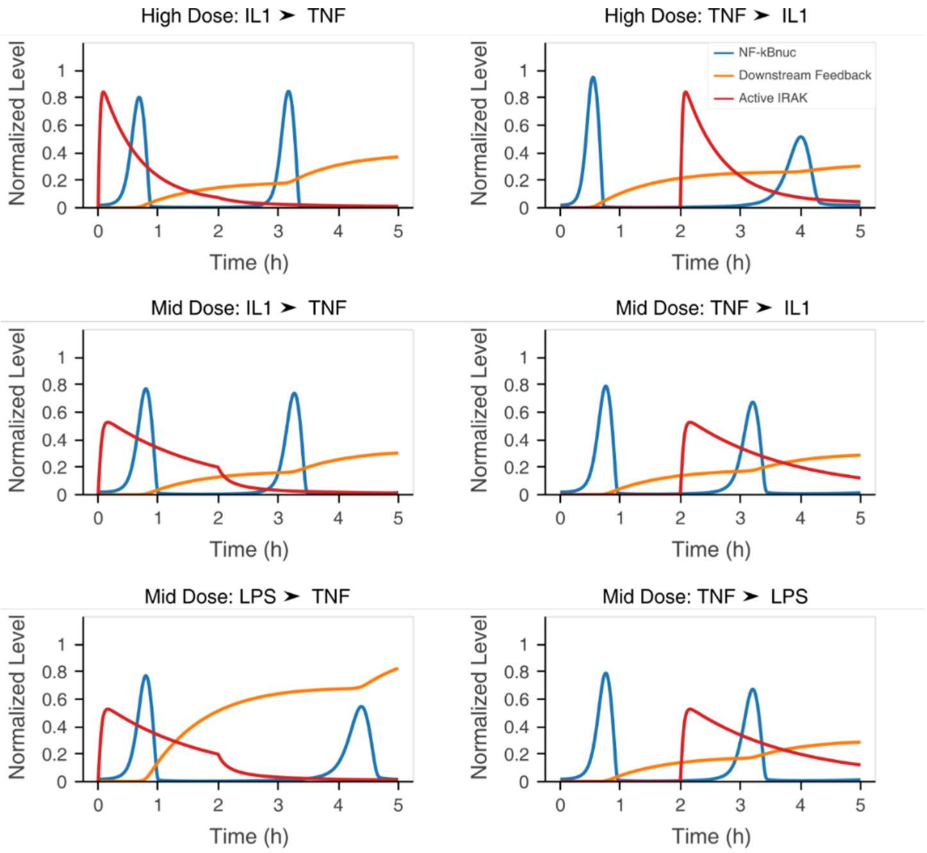


**Supplemental Figure 11:** Extension of Figure 5C–E. Each subplot shows the simulated NF-κB, active IRAK, and downstream feedback dynamics in response to the other sequences of stimuli not shown in Figure 5C–E. As in Figure 5C–E, the blue lines show the dynamics of nuclear NF- $\kappa$ B, the red lines for active IRAK1, and the orange lines show the dynamics for the downstream feedback component.

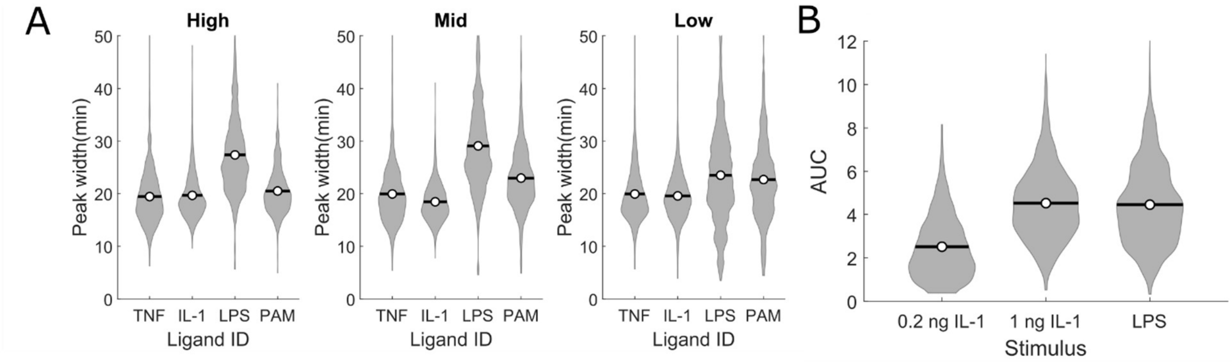


**Supplemental Figure 12:** A). Quantification of S1 peak width for each ligand from sequential stimulation data (S2-4). Width is defined as time-interval between half-maximal response on either side of the maximal response. B) Area under the curve (AUC) for the indicated stimulus responses. AUC calculated as trapezoidal approximation of integral over the stimulus interval.

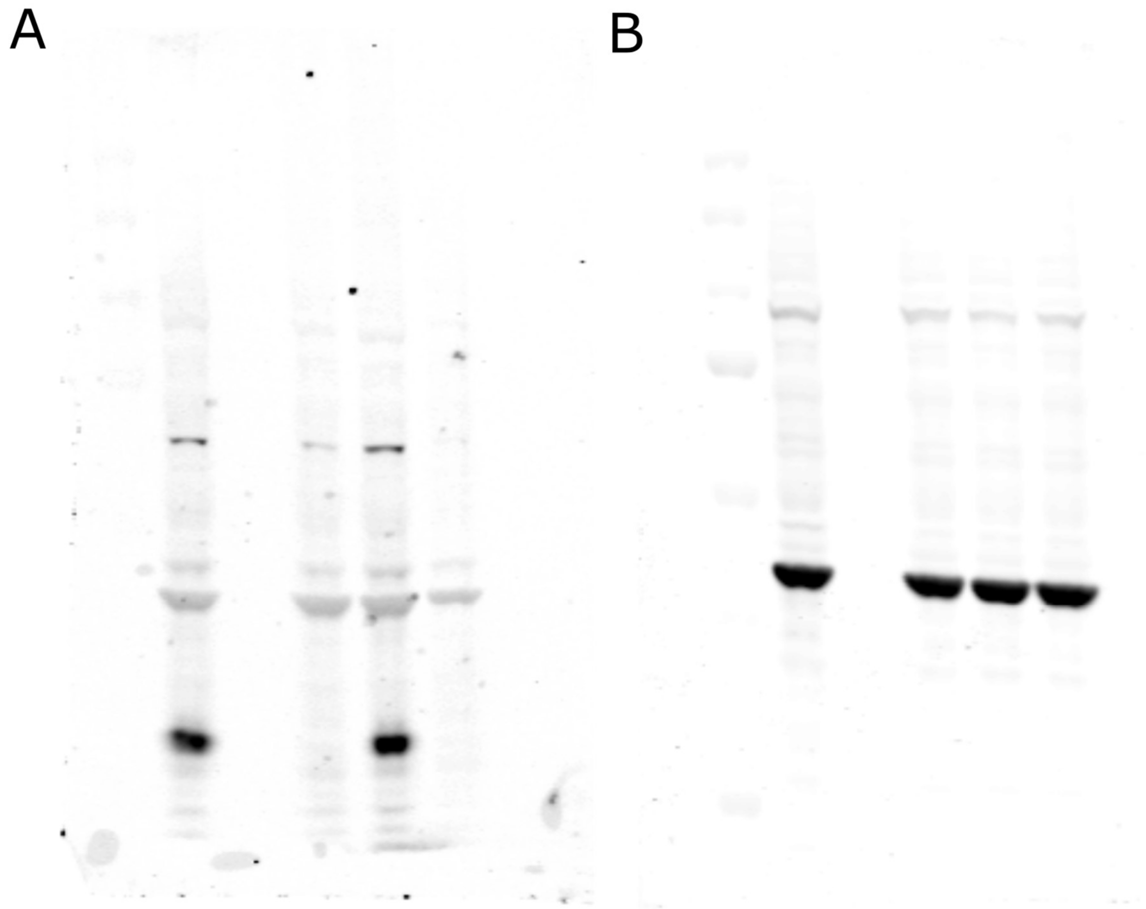


**Supplemental Figure 13:** Full western blots for MyD88 (A) and β-Actin (B).

**Supplemental Table 1:** Normalized log-counts-per-million (logCPM) for all 11,272 genes measured from sequencing data. LogCPM reported for three replicates of sequencing with IL-1β, LPS, and PAM treatment, as well as a no-treatment control.
